# Alternative splicing landscapes in *Arabidopsis thaliana* across tissues and stress conditions highlight major functional differences with animals

**DOI:** 10.1101/2020.11.10.374751

**Authors:** Guiomar Martín, Yamile Márquez, Federica Mantica, Paula Duque, Manuel Irimia

## Abstract

**Background:** Alternative splicing (AS) is a widespread regulatory mechanism in multicellular organisms. Numerous transcriptomic and single-gene studies in plants have investigated AS in response to specific conditions, especially environmental stress, unveiling substantial amounts of intron retention that modulate gene expression. However, a comprehensive study contrasting stress-response and tissue-specific AS patterns and directly comparing them with those of animal models is still missing.

**Results:** We generated a massive resource for *A. thaliana* (*PastDB*; pastdb.crg.eu), comprising AS and gene expression quantifications across tissues, development and environmental conditions, including abiotic and biotic stresses. Harmonized analysis of these datasets revealed that *A. thaliana* shows high levels of AS (similar to fruitflies) and that, compared to animals, disproportionately uses AS for stress responses. We identified core sets of genes regulated specifically by either AS or transcription upon stresses or among tissues, a regulatory specialization that was tightly mirrored by the genomic features of these genes. Unexpectedly, non-intron retention events, including exon skipping, were overrepresented across regulated AS sets in *A. thaliana*, being also largely involved in modulating gene expression through NMD and uORF inclusion.

**Conclusions:** Non-intron retention events have likely been functionally underrated in plants. AS constitutes a distinct regulatory layer controlling gene expression upon internal and external stimuli whose target genes and master regulators are hardwired at the genomic level to specifically undergo post-transcriptional regulation. Given the higher relevance of AS in the response to different stresses when compared to animals, this molecular hardwiring is likely required for a proper environmental response in *A. thaliana*.

## Background

Complex multicellular organisms generate a myriad of highly specialized cell and tissue types during their lifetime. Furthermore, they need to efficiently and rapidly react to a wide diversity of external stresses, both abiotic (e.g. high temperature or salinity) and biotic (e.g. bacterial infections), all requiring unique and highly coordinated molecular responses. Remarkably, all this is achieved by differentially regulating a single genome sequence to generate specific transcriptomes and proteomes for each environmental condition and cell or tissue type. In addition to the transcriptional control of gene expression, other less studied regulatory mechanisms are essential. Among them, alternative splicing (AS), the differential processing of introns and exons in pre-mRNAs to produce multiple transcript isoforms per gene, is the most important contributor to transcriptome diversification in both plants and animals [1–3]. The four major types of AS include exon skipping (ES), intron retention (IR) and alternative splice donor (ALTD) and acceptor (ALTA) choices, all of which have been observed in every major eukaryotic group, tracing the origin of these mechanisms back to the last common ancestor of eukaryotes [4]. However, the prevalence and proportions of each AS type varies widely across lineages [4–6].

AS can exert mainly two major molecular functions. On the one hand, it can expand proteome complexity by generating two or more distinct protein isoforms, which in some cases have been shown to present radically different functional properties [7]. On the other hand, by disrupting the main open reading frame (ORF) of the gene, AS can effectively lead to downregulation of its expression, either by creating truncated protein isoforms (which sometimes act as dominant negative [7]) and/or triggering non-sense mediated decay (NMD) [8]. Given the different proportions of AS types in each lineage, a higher relevance has been given to proteomic expansion associated with ES in animals [1], whereas in plants the focus has been put mainly into the regulation of gene expression by IR coupled to NMD [9–11]. The effects of ALTD and ALTA on gene function have not yet been thoroughly investigated in any species beyond a few special cases (e.g. NAGNAGs [12, 13]).

In contrast to the large number of studies exploring AS in animals, the prevalence, molecular functions and regulation of this mechanism in plants are much less studied. Several transcriptomic analyses have shown that AS is common in plants in general, and in *Arabidopsis thaliana*, in particular, detecting transcriptomic variation in up to 60-83% of multi-exonic genes [14–17]. These fractions are higher than those reported for fruitflies (20-37% [18]) but lower than those for humans, for which virtually every multi-exonic gene is alternatively spliced [19, 20]. These estimates, however, have been obtained using different methodologies and AS definitions, and thus cannot be readily compared. Moreover, although research on AS in plant systems has been gaining momentum in recent years, despite notable exceptions (e.g. [21]), most studies have so far focused on single genes or assessed the regulation of AS under specific environmental conditions, normally in response to an individual stress challenge [22]. This is in stark contrast with animals, for which most research has focused on tissue-specific regulation and on disease states [23, 24]. These differences in research focus in plants and animals may result in severe ascertainment biases in how we understand the roles and properties of AS in each lineage. Thus, a comprehensive study integrating harmonized AS profiles across multiple developmental time courses, mature cell and tissue types, as well as physiological and stressful environmental conditions in *A. thaliana* is still missing. Moreover, to properly evaluate if the widely assumed differences in the relative prevalence and molecular functions of AS between plants and animals are correct, a thorough comparison using equivalent approaches and datasets between both lineages is needed.

In this study, we have assembled a massive transcriptome-wide resource of quantitative profiles and associated genomic features for *A. thaliana* to address these gaps of knowledge. We characterized tissue-specific regulation and responses to different types of abiotic and biotic stress, and elucidated unique and common functional and regulatory properties of plant AS under these conditions. In addition, by comparing core sets of regulated AS events under each condition with those obtained using equivalent datasets from three animal models, we unveiled which properties are shared and which are lineage-specific. Finally, to boost AS research in plants, these transcriptomic quantifications and associated features are all made publicly available through *PastDB* (*Plant alternative splicing and transcription Data Base*; pastdb.crg.eu), the first web resource for *A. thaliana* integrating gene expression and AS profiles with multiple layers of genomic and regulatory information for a broad variety of samples.

## Results

### Transcriptome-wide landscapes of gene expression and alternative splicing in *A. thaliana*

We compiled a large panel of publicly available RNA-seq datasets for *A. thaliana* from the NCBI Short Read Archive (SRA). In total, we downloaded 516 individual SRA experiments from 63 independent studies, adding up to 20.4 billion reads (Additional file 1: Table S1). In most cases, to increase read depth for reliable AS quantification, we pooled together replicates from the same or similar experiments, using gene expression clustering of individual samples for verification of the groupings (see Methods for details). In total, we assembled a highly curated dataset comprising (Fig. 1): 90 samples covering multiple developmental time courses and mature cell or tissues types; 40 and 18 samples from abiotic and biotic stress experiments, respectively; 21 samples covering treatments with different light wavelengths; and 33 samples corresponding to experiments of 12 different mutants for genes involved in RNA processing.

**Figure 1.**
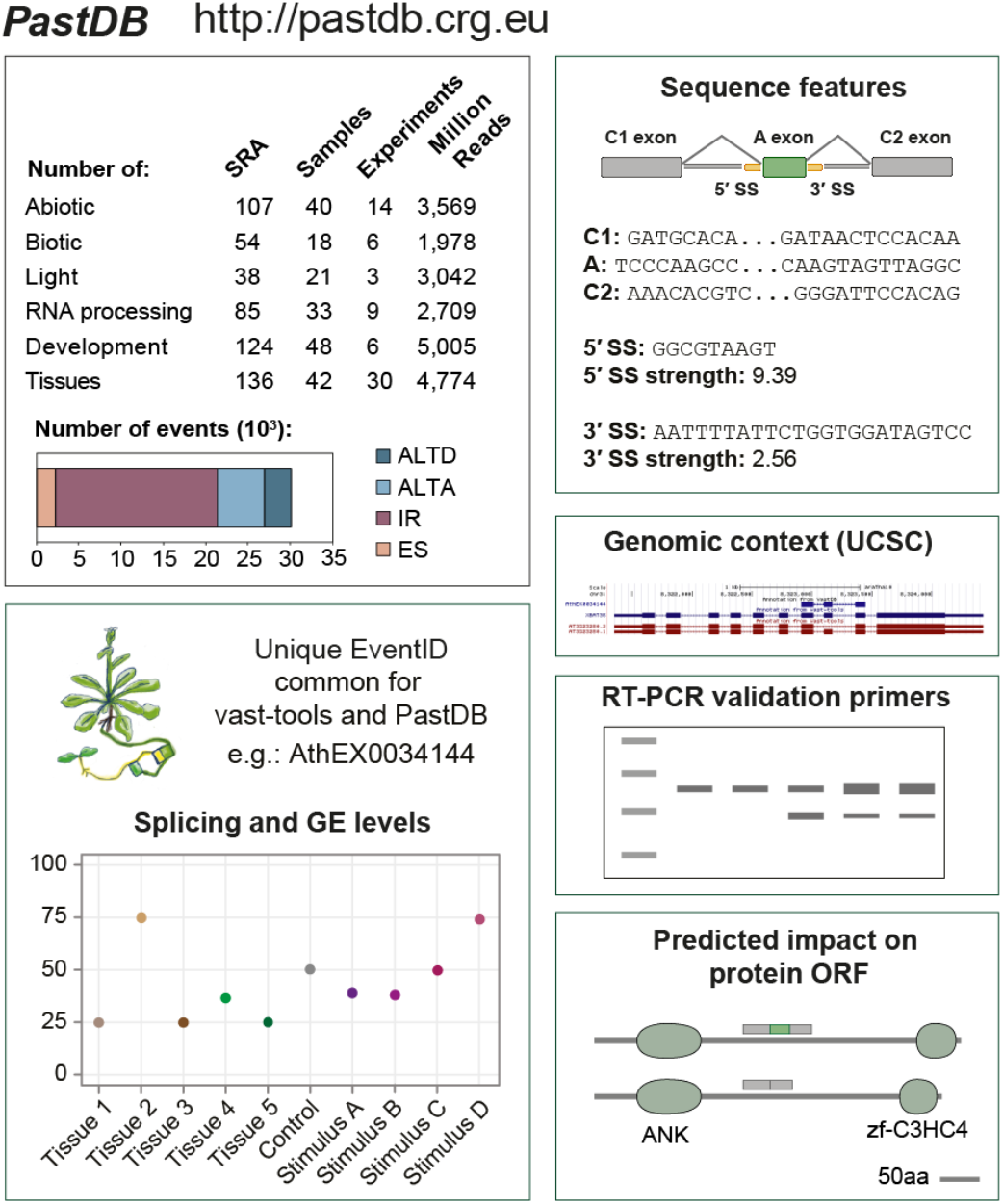
*PastDB*: An atlas of alternative splicing profiles and functional annotations in *A. thaliana*. *vast-tools* [25] was used to profile 516 independent RNA-seq datasets comprising a wide diversity of tissues, developmental stages, mutants for RNA-processing factors and physiological and stressful environments. The barplot shows the number of identified AS events by type across developmental time courses and mature cell and tissue types. Each AS event possesses a unique and constant event identifier (EventID) across *vast-tools* and *PastDB*, allowing profiling of new RNA-seq samples with *vast-tools* and direct contextualization of interesting events in *PastDB*. For each EventID (e.g. AthEX0034144), *PastDB* provides functional information, including a graphical display of splicing and gene expression levels across samples, sequence features, genomic context through the UCSC Genome Browser, suggested primers for RT-PCR validation, and multiple protein features (e.g., domains and disorder regions). AthEX0034144 corresponds to a previously described AS exon even located in *XBAT35* [75]. The number of AS events depicted corresponds to all *PastDB* samples excluding the RNA-processing factor experiments. ANK, ankyrin repeat; zf-C3HC4, RING finger domain.

We used the software *vast-tools* [25] to quantify steady-state mRNA levels (hereafter gene expression; GE) and alternative sequence inclusion levels for all four major AS types (ES, IR, ALTD and ALTA; see Methods). *vast-tools* has been widely used to detect and quantify AS in multiple vertebrate and non-vertebrate animals with a wide range of intron-exon structures, as well as unicellular eukaryotes, with very high validation rates [25–33]. Employing the framework of *VastDB* [25], we created *PastDB* (http://pastdb.crg.eu), the first web resource displaying comprehensive transcriptome-wide quantitative profiles for GE and AS in *A. thaliana* for a wide array of samples. In addition to RNA-seq based quantifications, *PastDB* also provides numerous associated genomic features for each AS event, including relevant sequences, splice site strength estimates, predicted impact on the main protein ORF, and suggestions of primers for RT-PCR validations (Fig. 1, Additional file 2: Figure S1).

Applying a strict definition for an event to be considered alternatively spliced (a percent inclusion level [PSI] between 10 and 90 in at least 10% of the samples with sufficient read coverage and/or a range of PSIs of at least 25) revealed thousands of *A. thaliana* AS events across the studied conditions (Fig. 1). We estimated that 62.2% of multi-exonic genes are alternatively spliced through at least one AS event across developmental and tissue samples, increasing up to 66.4% when stress and light experiments were also considered. Consistent with previous studies [4, 6, 16, 34], IR was the most prevalent type of AS genome-wide (Fig. 1). Furthermore, subsampling of increasingly larger numbers of samples showed that saturation has not been fully reached with >160 samples (Additional file 2: Figure S2). Using a less stringent definition of AS (i.e. a PSI between 5 and 95 in at least one sample; [35]), this percentage reached 94.8% of multi-exonic genes. On the other hand, around one third (33.7%) of multi-exonic genes harbored at least one AS event with extreme quantitative regulation (ΔPSI ≥ 50 between at least one pair of samples), producing distinct major transcript isoforms across conditions.

We also looked for PanAS events [25]: sequences that are alternatively spliced in the vast majority of sample types (i.e. with 10≤PSI≤90 in more than 80% of the samples with sufficient read coverage; see Methods for details). We identified 3,005 PanAS events, which were highly enriched for ALTA and ALTD events and strongly depleted for IR (Additional file 2: Figure S3a,b). PanAS events affected 2,295 genes, which corresponded to 14.2% of the multi-exonic genes that had the minimum read coverage required for this analysis (Additional file 2: Figure S3a, b), a fraction comparable to that of human multi-exonic genes with PanAS exons (18.5% [25]). As in mammals [25], genes harboring PanAS events were significantly enriched for DNA binding and transcriptional regulation. In addition, PanAS events in *A. thaliana* were enriched for RNA binding and splicing functions (Additional file 2: Figure S3c). Therefore, these results indicate that PanAS events are also modulators of regulatory hubs in plants.

### Identification of core genes regulated by AS and GE in response to internal and external stimuli

To begin assessing the functional roles of regulated AS in *A. thaliana*, we next characterized AS changes in response to different abiotic and biotic stresses. To standardize the comparisons, we selected ten independent RNA-seq experiments covering all major types of plant abiotic stress (drought, salt, heat, cold and ABA treatments) and biotic stress (bacterial and fungal infections), and performed individual pairwise comparisons between each control and treated sample (Additional file 3: Table S2; see Methods). These pairwise analyses identified 4,828 and 7,063 AS events that were differentially spliced (ΔPSI > 15) in at least one abiotic and biotic stress condition, respectively (Additional file 2: Figure S4a and Additional file 4: Table S3). IR generally increased upon abiotic stress induction, a widely reported trend [36], whereas exons more often showed decreased inclusion (Additional file 2: Figure S4a). With this information, we next defined a core set of 445 and 546 AS events in 368 and 453 genes that responded in a consistent manner (up- or down-regulating the inclusion of the alternative sequence) in at least two out of the five independent abiotic or biotic experiments, respectively (Fig. 2a, Additional file 2: Figures S4b,c and 5a and Additional file 4: Table S3; see Methods for details). We applied a similar approach for GE to define two core sets of 3,256 abiotic and 2,977 biotic transcriptionally responding genes (Additional file 2: Figure S6 and Additional file 5: Table S4). Gene Ontology (GO) analysis confirmed that these differentially expressed genes were strongly enriched for abiotic and biotic responsive genes (Additional file 2: Figure S7 and Additional file 6: Table S5). Furthermore, consistent with previous studies demonstrating a common transcriptional response to both types of stress [37, 38], the genes in both GE core sets were highly overlapping (∼40%). However, this common response was not observed at the post-transcriptional level, with <10% of events overlapping between the two stress AS core sets (Additional file 2: Figure S8).

**Figure 2.**
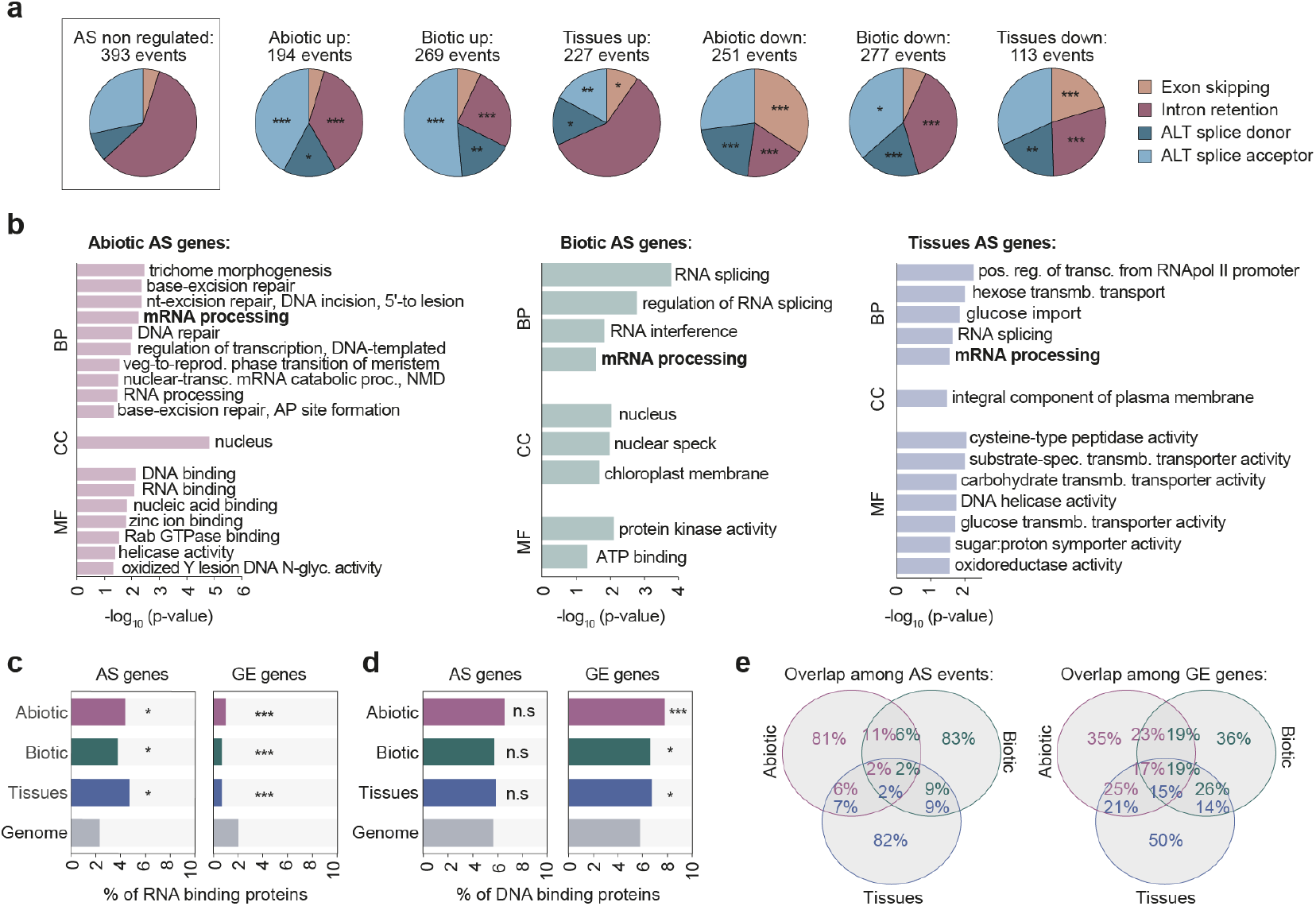
Identification and characterization of core sets of stress-responding and tissue-specific AS events. **a** Number of events by AS type for which the inclusion of the alternative sequence is differentially up- or down-regulated in each AS core set. **b** Enriched Gene Ontology categories of biological process (BP), cellular component (CC) and molecular function (MF) for genes from the abiotic (left), biotic (middle) and tissue (right) AS core sets. **c**,**d** Percentage of RNA (c) and DNA (d) binding proteins in the different AS and GE core sets. **e** Venn diagram showing the overlap among the three types of AS (left) or GE (right) core sets. *p*-values indicate statistical significance as assessed by two-sided Fisher’s Exact tests (a) or one-sided Fisher’s Exact tests (b-d). *P* < 0.05 (*); *P* < 0.01 (**); *P* < 0.001 (***). n.s., not significant.

Next, to define tissue-specific AS events, we compiled multiple independent RNA-seq experiments from six *A. thaliana* mature and developing samples (collectively referred as “tissue samples”), including inflorescences, leaves, roots, seeds, siliques and embryos (Additional file 2: Figure S9a and Additional file 3: Table S2). Following previous studies in animals [25, 27], we defined as tissues-specific those AS events with: (i) sufficient read coverage in at least two replicates of at least four different sample types, (ii) a |ΔPSI| ≥ 15 in the same direction between the target tissue and every other tissue, and (iii) an absolute average ΔPSI ≥ 25 (Additional file 2: Figure S5b; see Methods for details). In total, we found 330 AS events in 256 unique genes, which comprised the tissue-specific AS core set (Figure 2a, Additional file 2: Figure S9b and Additional file 4: Table S3). All AS types from the three core sets were similarly conserved at the genome level [39] across four Brassicaceae species, showing comparable or slightly higher levels than other alternative AS events and well above cryptic-like AS events (Additional file 2: Figure S10, see Methods for detail). In parallel, a GE core set of 5,480 tissue-specific genes was defined following a similar rationale (individual fold changes FC ≥ 3 in the same direction and an average FC ≥ 5; Additional file 2: Figure S9c and Additional file 5: Table S4).

As a control group for the three core sets of AS events (abiotic stress, biotic stress and tissue-specific), we also identified a subset of events whose inclusion does not change substantially in response to any abiotic or biotic stress or across tissue samples (non-regulated AS events, “AS-NR”; see Methods for details). Consistent with the general AS pattern in *A. thaliana* (Fig. 1) and previous reports [6, 16, 34], IR was the most prevalent type in this control set (Fig. 2a). Interestingly, however, IR events were significantly reduced in the three core sets of regulated AS in favor of the other types of AS events (4.16e-21 ≤ *P* ≤ 0.016 for each of the three sets respect to AS-NR events, Fisher’s Exact test). Together with the findings for PanAS events, this result indicates that the relative contribution of IR to regulated transcriptome remodeling in *A. thaliana* might be lower than commonly assumed.

The three sets of genes regulated by AS (AS core sets) were enriched for mRNA processing and related functions (Fig. 2b), which were also observed for each AS event type separately (Additional file 2: Figure S11). In particular, RNA binding proteins (RBPs) were significantly enriched among the three AS core sets (Fig. 2c; 0.012 ≤ *P* ≤ 0.036, Fisher’s Exact test), whereas no bias was observed for transcription factors (TFs) and other DNA binding proteins (Fig. 2d). In contrast, transcriptionally regulated genes from the three categories (GE core sets) were significantly enriched for TFs and depleted for RBPs (Fig. 2c, d; 1.73e-5 ≤ *P* ≤ 0.046, Fisher’s Exact test). A similar specialization of both molecular mechanisms impacting regulatory hubs had been previously shown in early response to light stimuli [40], but not under stress conditions or across tissue types, suggesting that this regulatory specialization is widespread. Remarkably, these differential enrichments were tightly mirrored at the genomic level: whereas genes encoding RBPs had weaker splice sites and shorter promoters with fewer TF binding sites compared to the genome average, genes encoding DNA binding proteins displayed the exact opposite patterns (Additional file 2: Figure S12 and see below).

In addition to the shared functions of genes from the AS core sets in the regulation RNA-related processes, each set showed specifically enriched GO terms, including DNA repair (abiotic stress), protein kinases (biotic stress) or membrane transporters (tissues) (Fig. 2b). We also observed distinct enrichment of subcellular locations for each subset of proteins with AS regulation (Additional file 2: Figure S13). Further supporting a substantial degree of differentiation of the effector genes regulated by AS in different conditions, the overlap of the three AS sets is very small compared to that of the transcriptionally regulated genes, both at the AS event (Fig. 2e) and gene (Additional file 2: Figure S14) levels.

### Molecular functions of AS regulated genes in *A. thaliana*

To evaluate the functional relevance of these regulated AS events, we next investigated their predicted impact on the canonical ORF. Remarkably, for regulated ES and ALTA/ALTD events, we observed a depletion in the proportion of cases predicted to generate alternative protein isoforms in favor of alternative sequences with regulatory potential (6.49e-8 ≤ *P* ≤0.048, Fisher’s Exact test; Fig. 3a). These include AS events located in the coding region that will likely alter transcript and/or protein levels (e.g. by excluding/introducing a premature termination codon leading to NMD) and sequences in the untranslated regions (UTRs), which may also have regulatory consequences (e.g. affecting transcript stability or translation efficiency; [41]). While it is well known that IR can lead to ORF disruption and NMD-mediated degradation [9, 10, 28, 42, 43], the contribution of other AS types to the modulation of functional protein levels is less established. Therefore, to test this hypothesis for other types of regulated AS, we first compared the expression levels of genes containing regulated EX, ALTA or ALTD events in WT and *upf13* double mutants, which show impaired NMD [10]. Strikingly, these genes were significantly upregulated in *upf13* seedlings, indicating that a substantial fraction of their transcriptomic output is targeted to NMD (Fig. 3b, Additional file 2: Figure S15a). As expected, the same pattern was observed for genes with regulated IR events (Fig. 3b, Additional file 2: Figure S15a).

**Figure 3.**
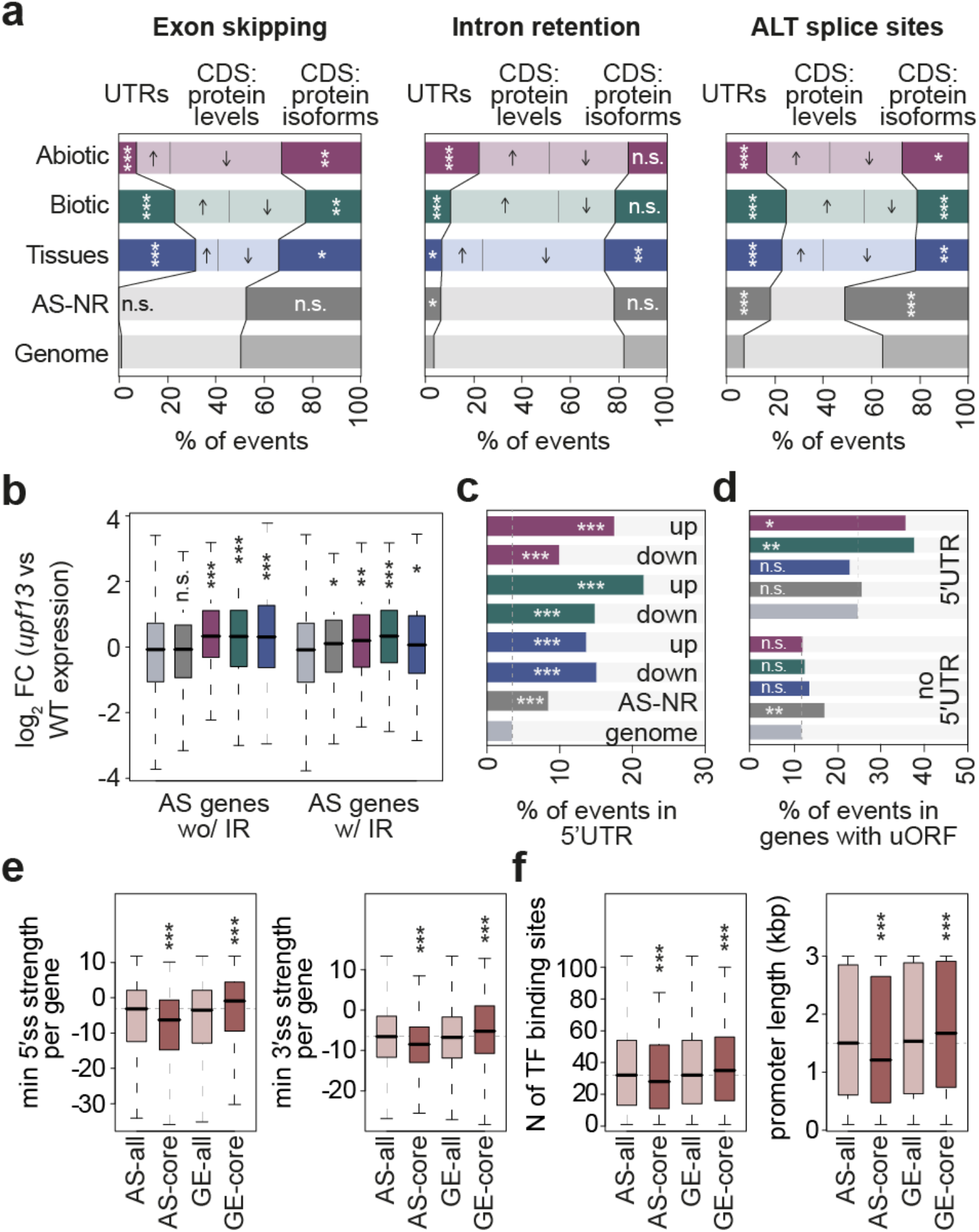
Functional impact and genomic features of differentially spliced events in *A. thaliana*. **a** Percentage of exon skipping (left), intron retention (middle) and alternative splice donor and acceptor sites (ALT splice sites, right) of events belonging to the different AS core sets located in UTRs or CDS gene regions. Among the latter category, the percentage of events that potentially affect protein levels or produce alternative protein isoforms is indicated (see Methods for details). Arrows indicate whether the levels of proteins are expected to increase or decrease in the stress or tissue-specific condition. **b** Log2 of the fold change in the expression values of *upf1/upf3* double mutant (*upf13*) compared to WT for genes belonging to the different AS core sets. Data from [10]. **c** Percentage of events for which the inclusion of the alternative sequence is up- or down-regulated from each AS core set that occur in the 5′ UTR of their respective genes. **a-c** “Genome” contains all multi-exonic *A. thaliana* genes fulfilling the coverage criteria used for the corresponding AS analysis. **d** Percentage of events from each subcategory that occur in genes with uORFs. Data from [76]. **e** Distributions of the lowest 5′ splice site (left) or 3′ splice site (right) strength per gene for each AS and GE core set. Values correspond to the splice site with the lowest maximum entropy score in each gene, calculated as in [77] (see Methods). **f** Number of binding sites of *A. thaliana* transcription factors (TF) per gene’s promoter (left) and promoter length for each gene (right). Promoter information was obtained from [78]. **e-f** AS-all and GE-all sets contain genes fulfilling the coverage criteria used for GE and AS analyses (see Methods for details). *p*-values indicate statistical significance as assessed by two-sided Fisher’s Exact tests (a), one-sided Fisher’s Exact tests (c,d) or Wilcoxon Rank-Sum test (b,e). *P* < 0.05 (*); *P* < 0.01 (**); *P* < 0.001 (***). AS-NR, alternative spliced-non regulated; n.s., not significant; w/, with; wo/, without.

Because of the strong enrichment of regulated AS events located in the UTRs, we next studied this pattern in more detail. The enrichment was observed for 5′ UTRs in all AS core sets (Fig. 3c; 1.26e-29 ≤ *P* ≤ 1.7e-6, Fisher’s Exact test) and in all AS event types (Additional file 2: Figure S15b), but not for 3′ UTRs (Additional file 2: Figure S15c). Moreover, it was particularly strong for those cases in which the inclusion of the alternative sequences was upregulated in response to biotic and abiotic stimuli, indicating that transcripts from these genes tend to have longer 5′ UTRs in stress conditions. Remarkably, stress-regulated AS events located in the 5′ UTR were specifically enriched in genes with upstream ORFs (uORFs) (6.7e-3 ≤ *P* ≤ 0.049, Fisher’s Exact test; Fig. 3d). These observations suggest a regulatory model by which these stress-specific longer 5′ UTRs introduce uORFs impacting translation in response to environmental challenges. Moreover, we found that transcripts from genes regulated by AS under stress conditions (yet not across tissue types) had a significantly lower half-life than other gene sets, consistent with a fast turn over that might be expected for stress-response genes (Additional file 2: Figure S15d; half-life data from [44]).

Overall, these results indicate that events from all regulated AS core sets play an important regulatory role in the control of total transcript and protein levels under a wide range of physiological and stress conditions. Interestingly, despite this shared molecular function, the overlap between genes from the AS and GE core sets was minimal (Additional file 2: Figure S16). Hence, we hypothesized that AS-regulated genes rely inherently more on AS and less on transcription for the regulation of their functional mRNA levels, whereas GE-regulated genes are less prone to post-transcriptional control, and that these differences must be mirrored at the genomic level. In agreement with this, we found that, compared to the genome average, genes from the AS core sets had (i) significantly weaker 5′ and 3′ splice sites, (ii) longer gene bodies with higher numbers of introns, and (iii) shorter promoter regions with fewer TF binding (Fig. 3e, Additional file 2: Figure S17a-f). On the other hand, genes from the GE core sets showed the exact opposite pattern, with stronger splice sites, smaller gene bodies with fewer introns and longer promoters with more TF binding sites. Thus, these findings suggest that genes regulated by AS or GE are globally hardwired at the genomic level to be controlled mainly by only one of the two regulatory layers. Remarkably, given that RBPs and TFs show a similarly contrasting pattern (Additional file 2: Figure S12), our results indicate that this hardwiring also affects the master regulators of the two layers, which we also showed to be more likely to exhibit self-regulation and avoid inter-layer cross-regulation (Figure 2c, d).

### Regulatory insights into AS core set events in *A. thaliana*

To investigate the genomic features associated with AS regulation in more detail, we first analyzed events from each AS type separately using *Matt* [45], which extracts and compares over 50 features associated with AS regulation (see Methods). In general, up- and down-regulated events from each regulated AS core set (abiotic stress, biotic stress, and tissue-specific) showed similar genomic features (Additional file 2: Figures S18a-S21a). For simplicity, we thus reanalyzed the three categories together grouped by AS type and by whether the inclusion of the alternative sequence was up- or down-regulated in response to stress or specifically in a given tissue (Fig. 4a-d, Additional file 2: Figures S18b-S21b). Several clear patterns emerged. First, both up- and down-regulated ES events showed the main hallmarks of exon definition, i.e. shorter exons flanked by longer introns (Fig. 4a), despite the fact that plant splicing is largely dominated by intron definition [4, 46, 47]. Furthermore, exons whose inclusion was down-regulated were flanked by introns with higher GC content, whereas the exons themselves showed lower GC content for up-regulated ES events. Another major difference between both exon sets was that down-regulated exons harbored the weakest splicing signals, including both 5′ and 3′ splice sites and a larger distance between the branch point and the 3′ splice site (BP-AG distance).

**Figure 4.**
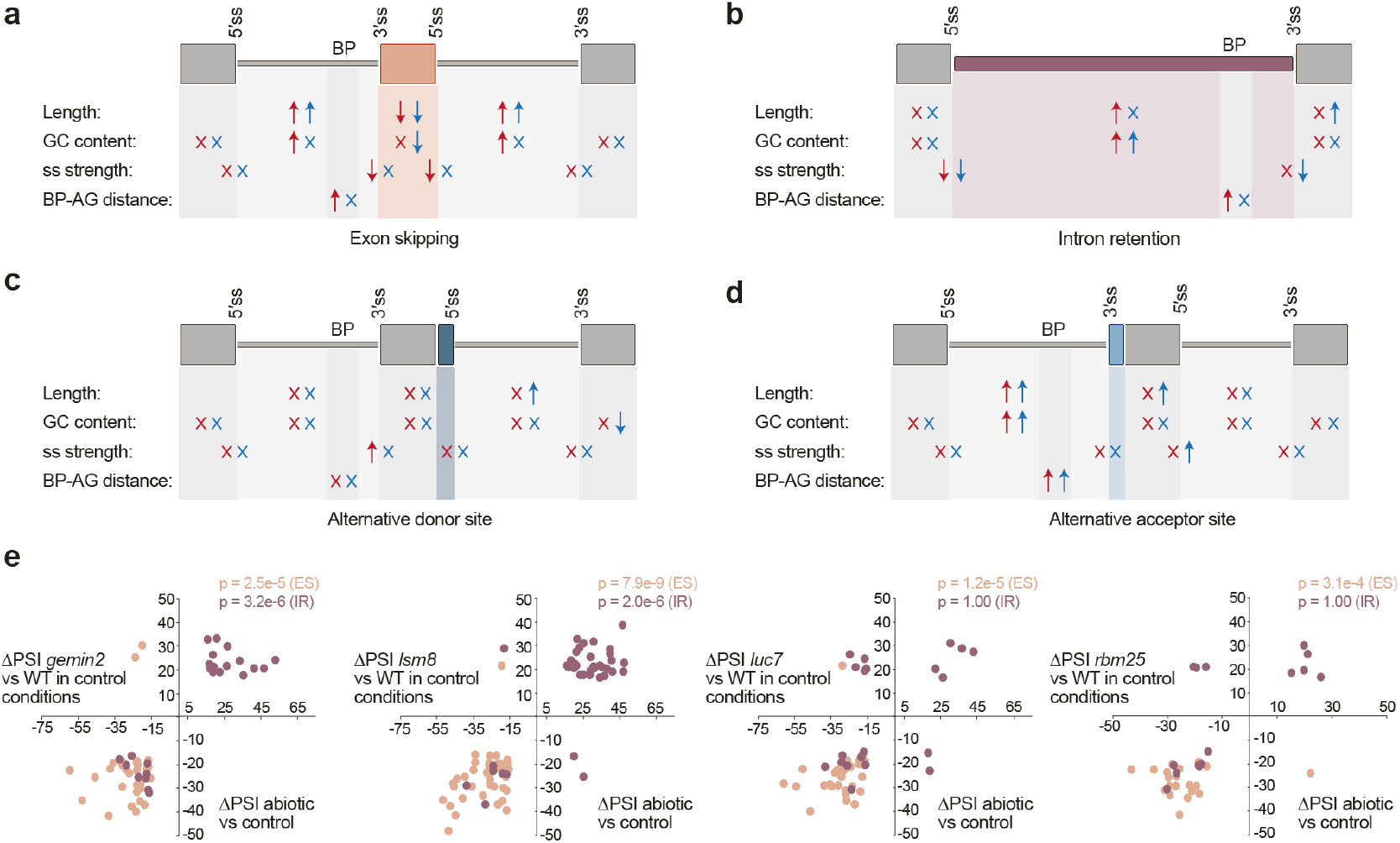
Schematic representation of genomic regulatory features associated with differentially spliced AS events. For exons (**a**), introns (**b**), alternative donor (**c**) and acceptor (**d**) sites belonging to any of the three AS core sets, arrows summarize which features show statistically significant differences with respect to the Genome set of events and the direction of these differences (higher or lower). “X” indicates no statistically significant difference. Features include: length of the alternatively spliced sequence together with its downstream and upstream exons/introns, GC content, maximum entropy (MaxEnt) score for 5′ and 3′ splice sites (ss strength), and distance between the branch point and the 3′ splice site (BP-AG distance). Differences are indicated separately for events for which the inclusion of the alternative sequence is down- (red) and up- (blue) regulated in the core sets. The genome set contains events fulfilling the read coverage criteria used for AS analyses. Genomic features were extracted using *Matt* [45]. ss, splice site. **e** Scatterplot of ΔPSI values for IR and ES events belonging to the abiotic stress AS core set whose inclusion is affected (|ΔPSI|>15) by the studied splicing factors. ΔPSI between abiotic and control conditions corresponds to the average ΔPSI of experiments in which the event was found to change significantly. *p*-values correspond to Binomial tests for quadrants I and III vs. the total.

In the case of IR events, introns whose inclusion was up- and down-regulated presented a mixture of shared and unique features (Fig. 4b). Whereas both groups of introns (as well as NR-AS IR events; Additional file 2: Figure S19b) had higher GC content and weaker 5′ splice sites, only up-regulated introns showed weaker 3′ splice sites, and only down-regulated IR events were longer with increased BP-AG distances. The increased intron length is in stark contrast to what has been reported for differentially regulated IR events in vertebrates, which were significantly shorter than other introns [28, 48]. Finally, ALTA and ALTD events were associated with longer upstream and downstream introns, respectively; however, unexpectedly, the actual alternative splice sites were not weaker than those of the control sets (Fig. 4c, d).

These analyses revealed an association of general genomic features with specific types of regulated AS, suggesting that modulation of the core splicing mechanism may underlie some of the observed regulatory patterns. In line with this, exploration of the available RNA-seq data for mutants of various RNA processing factors (Additional file 7: Table S6) revealed that depletion of some core spliceosomal components (*luc7, gemin2, lsm8* and *rbm25*) had a strong effect specifically on ES and IR events regulated by abiotic stress conditions (Fig. 4e, Additional file 8: Table S7). In particular, loss-of-function mutations in *gemin2* and *lsm8*, which are important for multiple steps of spliceosomal assembly [49, 50], caused both increased IR and ES (Additional file 2: Figure S22), with these changes tightly mimicking the AS patterns observed under abiotic stress (7.8e-9 ≤ *P* ≤ 2.5e-5, Binomial test; Fig. 4e, Additional file 8: Table S7). In the case of *luc7* and *rbm25*, which are specifically associated with the U1 snRNP and 5′ splice site recognition [51, 52], their disruption caused mainly ES (Additional file 2: Figure S22), likely by compromising exon definition. Strikingly, these shifts tightly reproduced the changes observed for ES in abiotic stress (1.2e-5 ≤ *P* ≤ 3.1e-4, Binomial test; Fig. 4e, Additional file 8: Table S7). Overall, these mutants produced consistent alterations in 49.6% and 64.2% of abiotic IR and ES events, respectively, suggesting that AS changes observed under abiotic stress may be caused, at least in part, by a disruption in core spliceosomal activity that specifically affects a subset of AS events with unique genomic features, including generally weaker splicing signals. This is consistent with previous studies on the effect of high temperature on AS in animals [53, 54] and the specific increase in IR and ES, both hallmarks of altered spliceosomal activity, observed upon plant exposure to different abiotic stresses (Additional file 2: Figure S4 and [36]). In addition, we also found that *GRP7* overexpression caused down-regulation of ES events from the abiotic AS core. Finally, it is worth noting that none of the studied RBPs appears to be significantly associated with the AS events of biotic stress and tissue core sets (Additional file 8: Table S7), suggesting that their patterns may be controlled by more specific RBPs yet to be characterized.

### Functional comparative analyses between plant and animal AS-regulated genes

Our results thus far highlight multiple patterns for AS events regulated in stress conditions and across tissues in *A. thaliana*. To put them into a broader evolutionary context, we compared them with equivalent regulated AS event sets from three major animal models — *Caenorhabditis elegans, Drosophila melanogaster* and *Homo sapiens* — with widely divergent genomic structures, including intron density and length, which are major determinants of AS regulation (Additional file 2: Figure S23). For this purpose, we collected RNA-seq data for five individual abiotic and biotic experiments for each species and multiple individual samples for five differentiated tissues (Additional file 9: Table S8), which were all processed using *vast-tools*. First, we estimated the levels of AS in each species using these comparable sets of samples. For each species, we subsampled an increasingly larger number of samples and scored the proportion of multi-exonic genes with at least one AS event with inclusion levels between 10 and 90 (or 20-80 or 30-70) in at least one sample. Considering all AS types together, human had the highest fraction of alternatively spliced genes, followed by *A. thaliana* and *D. melanogaster* with comparable AS levels, which were substantially higher than those of *C. elegans* (Additional file 2: Figure S24a). Interestingly, when each AS type was analyzed separately, *A. thaliana* showed particularly elevated levels of ALTA and, as expected, relatively reduced ES compared with animal species (Additional file 2: Figure S24b).

Next, we defined core sets of regulated AS events for each animal species following the same definitions that we used for *A. thaliana* (Additional file 2: Figure S5; see Methods for details). These regulated sets ranged from 67 AS events in 61 genes for biotic stress in *D. melanogaster* to 3,076 in 1,935 genes for tissues in *H. sapiens* (Additional file 2: Figure S25 and Additional file 10: Table S9). These core sets of AS-regulated genes showed distinct functional features in animals and *A. thaliana* based on comparative GO analyses (Fig. 2b, Additional file 2: Figures S26-28 and Additional file 11: Table S10). For instance, consistent with previous reports [25, 55], genes with tissue-specific AS were enriched in cytoskeleton and protein binding GO terms in all studied metazoans, but these terms were not observed for *A. thaliana*. Similarly, functions associated with biotic stress included transmembrane receptors or transporters in all animals but not in *A. thaliana*, for which such terms were instead enriched among the tissue-specific AS core set. Moreover, the enrichment of mRNA processing-related categories in all *A. thaliana* sets was not readily observed in the three animal organisms. Consistent with these differences, there was very little overlap between the AS core sets from *A. thaliana* and those of animals at the orthologous gene level, and none at the AS event level (Additional file 2: Figure S29, see Methods for details).

Interestingly, while the size of the three core sets of regulated AS events was largely comparable in *A. thaliana*, the three animal species had a striking overrepresentation of tissue-specific AS events (Fig. 5a). These differences could also be observed in the range of PSIs across experiments (Fig. 5b). Thus, these results suggest that the contribution of regulated AS to establish tissue-specific transcriptomic differences in animals is quantitatively much greater than that of the response to stresses. To specifically evaluate this hypothesis, we calculated the relative contribution of each regulatory axis (abiotic stress, biotic stress and tissues) to the total PSI variation for each AS event genome-wide (see Methods). When comparing the relative PSI variation of each stress type versus tissues in *A. thaliana*, we observed a similar contribution for tissues and stress, even slightly skewed towards stress for biotic experiments (Fig. 5c). Strikingly, the distributions were highly skewed towards tissues for the three animal species (Fig. 5c), even when the tissues with the strongest AS patterns are excluded (i.e. neural, muscle and testis; Additional file 2: Figure S30).

**Figure 5.**
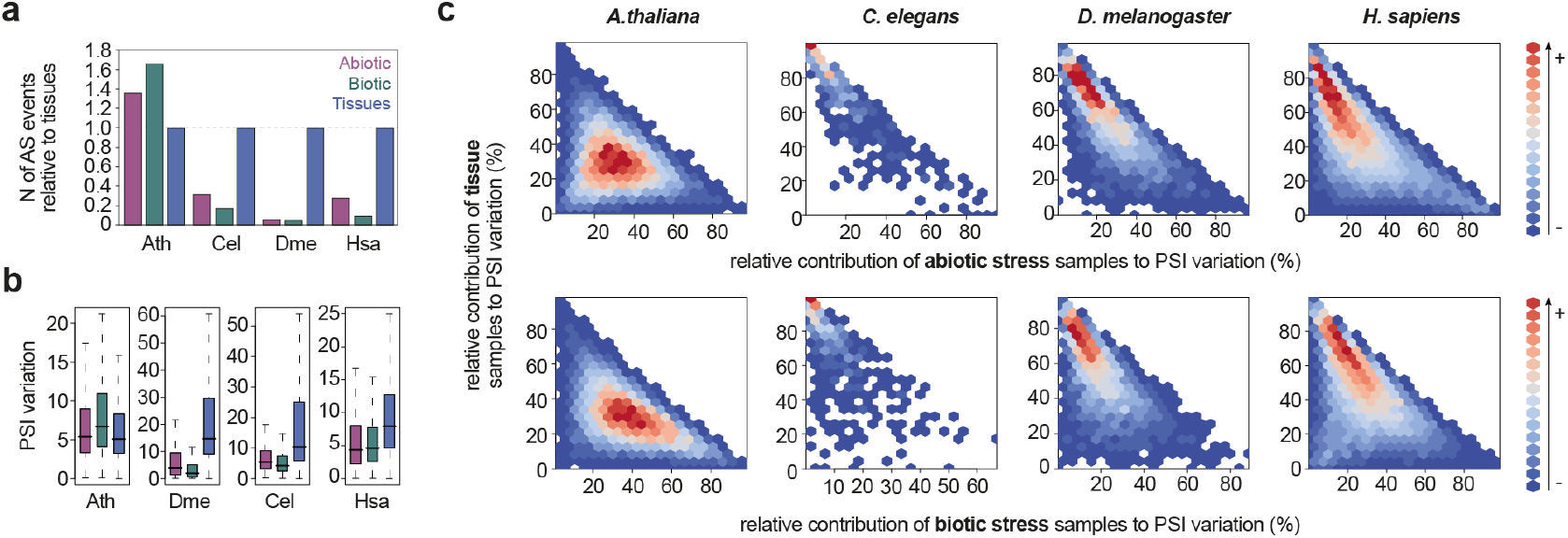
Relative contribution of tissue and stress samples to PSI variation in different organisms. **a** Number of total events in each AS core set relative to the tissue AS core set in each species. **b** Distribution values of the maximum ΔPSI among abiotic and biotic experiments and the PSI range across tissues for each species. **c** Comparison of the relative contribution to the total PSI variation of the tissue samples vs. that of the abiotic (up) or biotic (bottom) stress experiments in each species. The total PSI variation for each AS event is calculated as the sum of the three relative contributions: (i) the PSI range across tissues, (ii) the maximum ΔPSI among the abiotic stress experiments, and (iii) the maximum ΔPSI among the biotic stress experiments (see Methods for details). The color code represents the number of AS events found in each possible intersection between the relative contributions (in percentage) for each set of samples. Ath, *A. thaliana*; Cel, *C. elegans*; Dme, *D. melanogaster*; Hsa, *H. sapiens*.

Animals and *A. thaliana* also differed in the proportion of AS types within each regulated AS set, with higher fractions of ES events in metazoans (Fig. 6a). However, the relative proportions among regulated AS sets bore certain similarities across species. For example, ALTA and ALTD were over-represented under both biotic and abiotic stress conditions in all species. On the other hand, the highest fractions of IR were unexpectedly observed for the tissue-specific sets. Next, we analyzed the impact of animal regulated AS in the main ORF. In stark contrast to *A. thaliana*, the three animals showed a strong enrichment for protein isoforms (Fig. 6b), a pattern that was particularly prominent for tissue-specific ES and ALTA/ALTD events and was also observed for AS-NR events in most cases. In line with this finding, analyses of RNA-seq data from NMD-depleted cells from *C. elegans, D. melanogaster* and *H. sapiens* showed that AS-regulated genes in these cells exhibited no significant differences in mRNA steady state levels respect to controls (Additional file 2: Figure S31), unlike the equivalent AS gene sets in *A. thaliana* (Fig. 3b). However, similar to plants, we observed a significant enrichment for AS events in the UTRs for *D. melanogaster* and *H. sapiens*, particularly for stress-regulated AS events (Fig. 6b), consistent with previous analyses [56] and suggesting that regulatory roles for stress-regulated AS have evolved in both plant and animal species.

**Figure 6.**
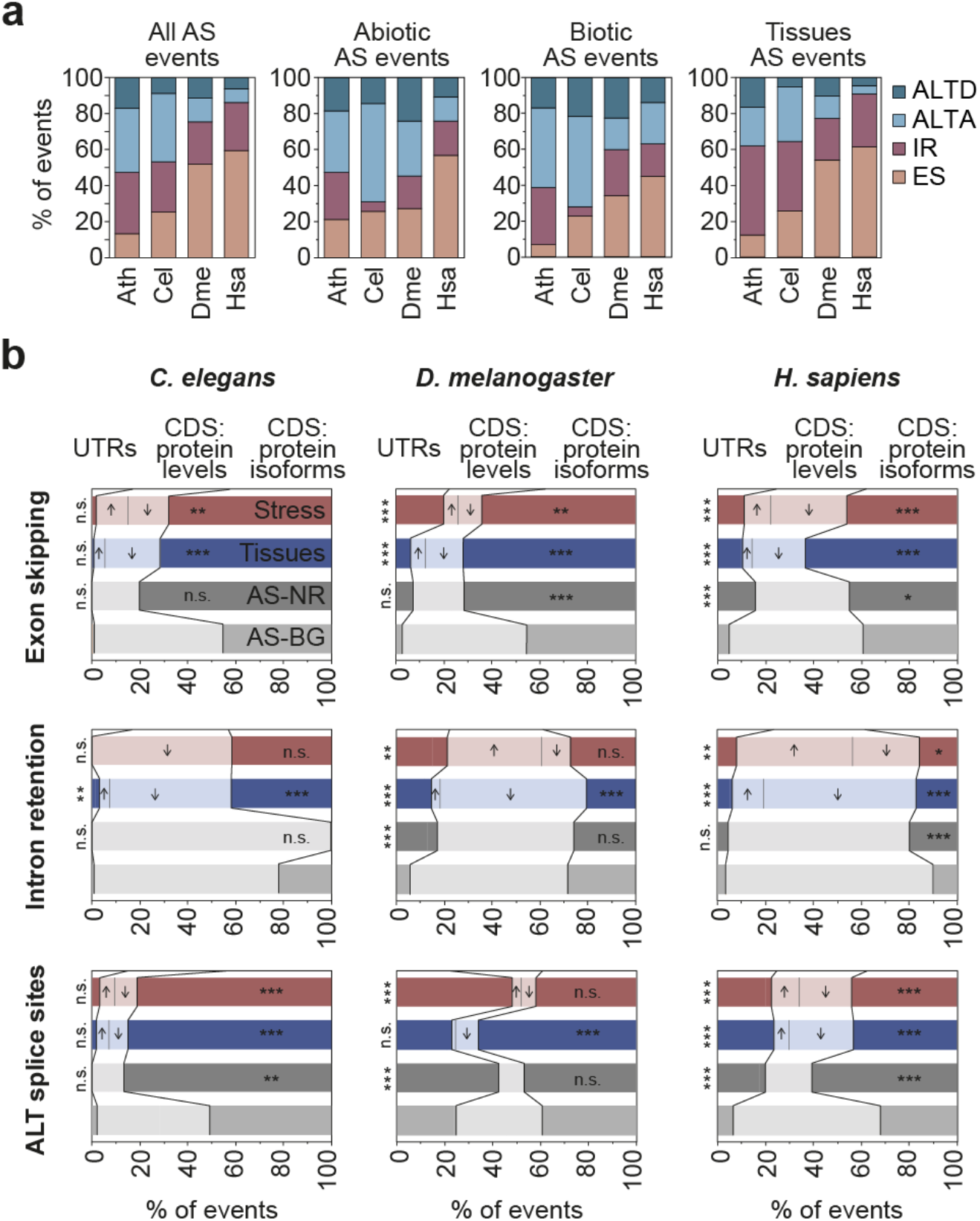
Functional impact of differentially spliced events in *C. elegans, D. melanogaster* and *H. sapiens*. **a** Proportion of each type of AS event in the three AS core sets for each studied species. **b** Percentage of exon skipping (top), intron retention (middle) and alternative splice donor and acceptor sites (ALT splice sites, bottom) events belonging to the different AS core sets located in UTRs or CDS gene regions. Among the latter category, the percentage of events that potentially affect protein levels or generate alternative protein isoforms is indicated (see Methods for details). Arrows indicate whether the levels of proteins are expected to increase or decrease in the stress or tissue-specific condition. Given the low number of events in some types of Abiotic and Biotic AS core sets they are joined in a single group: Stress core set. *p*-values indicate significance as assessed by Wilcoxon Rank-Sum tests (P < 0.05 (*); P < 0.01 (**); P < 0.001 (***)). AS-NR, alternative spliced-non regulated; n.s., not significant. Ath, *A. thaliana*; Cel, *C. elegans*; Dme, *D. melanogaster*; Hsa, *H. sapiens*.

## Discussion

This study reports the most comprehensive AS analysis conducted so far in *A. thaliana*, focusing both on stress responses as well as on tissue-specific patterns, the latter of which has been largely understudied in plants. Moreover, we performed the first direct systematic comparison between plant and animal AS using equivalent RNA-seq datasets. Through these analyses, we shed light on the prevalence, regulation, molecular functions and evolutionary aspects of AS in *A. thaliana*.

We found high levels of AS in *A. thaliana*, comparable to those of a complex metazoan such as *D. melanogaster*. Although these fractions ultimately depend on the exact definition of AS and the number and type of samples used, we estimate that around ∼70% of *A. thaliana* multi-exonic genes are at least moderately alternatively spliced and that more than one third show evidence for strong alternative processing. Interestingly, although we found that, in line with previous studies [4, 6, 16, 34], IR is the most common AS type genome-wide, we showed that the other AS types (ES, ALTA and ALTD) are in fact overrepresented compared to IR in most of the categories of regulated — and thus more likely functional — AS we have identified. These include AS events differentially regulated upon multiple stresses, among tissues, or that are broadly alternatively spliced across conditions (PanAS events). Moreover, in comparison with metazoans, ALTA events are particularly prevalent in *A. thaliana*. Our results therefore indicate that the functional importance of non-IR AS to the dynamic remodeling of plant transcriptomes may have so far been underrated.

A central goal of our study was to elucidate how plants utilize AS in response to different external and internal stimuli. We identified largely non-overlapping core sets of genes that are differentially alternatively spliced in response to each type of stress or in specific tissues. By combining different approaches and data sources, we determined that most of these tightly regulated AS events are likely to play a role modulating the final levels of functional protein products. Such a role has been reported in multiple previous studies in association with IR [57]. Here, we established it for IR but, importantly, also for the other AS types. We showed that regulated events from all AS types that fall in the coding sequence often disrupt the main ORF and that the associated genes are actively targeted by NMD. Moreover, we identified a novel signature that strongly affects the 5′ UTRs and the presence of uORFs. Given that the majority of these events lead to sequence inclusion in the 5′ UTRs upon stress, we speculate that they introduce stress-specific uORFs that impact the translation of the host genes in response to challenging environments. This and alternative functional hypotheses should be evaluated in future studies.

Altogether, these results thus place AS in *A. thaliana* as an extra regulatory layer for the control of gene expression in response to stimuli. Importantly, our data indicate that this layer is not redundant with that of transcriptional regulation (e.g. by serving as a secondary safety mechanism), but is largely non-overlapping and complementary. Consistent with other studies [40, 58–63], we showed that the majority of genes regulated by AS and GE under different conditions do not overlap. Remarkably, we found that the propensity for a gene to be regulated by either AS or GE seems to be hardwired in the genome. Genes regulated by AS have particularly weak splice sites, favoring post-transcriptional processing, and, at the same time, are associated with shorter promoters with fewer TF binding sites than the genome average, implying poor transcriptional regulatory potential. The pattern is the exact opposite for GE-regulated loci, with more complex promoters to enable transcriptional regulation, and strong splice sites, which should facilitate efficient constitutive splicing. Strikingly, these genomic patterns are also observed for the master regulators of each layer: while RBPs have the signature of AS-regulated genes, TFs and other DNA binding proteins exhibit that of GE-regulated genes, and this is consistent with their differential regulation by each mechanism. Therefore, taken together, our findings suggest that *A. thaliana* has established two largely parallel and specialized regulatory layers to efficiently tackle challenging environmental conditions. Why these independent layers may be required for an efficient response remains to be established. However, as each mechanism has its own temporal dynamics and variation range, it is certainly possible that post-transcriptional regulation may provide some adaptive advantages to certain genes that cannot be fulfilled through transcriptional regulation and vice-versa.

Our study also provides the first direct comparison between regulated AS in *A. thaliana* and animals. This comparison revealed major qualitative and quantitative differences in how each lineage employs this post-transcriptional regulatory layer. First, whereas much of *A. thaliana*’s AS is regulated in response to stress conditions, the vast majority of animal AS is controlled in a tissue-dependent manner. Moreover, as mentioned above, *A. thaliana*’s AS mainly impacts functional mRNA and protein production, both in response to stress and across tissues. By contrast, in the case of animals, although we also found evidence for a subset of AS events with regulatory potential, there is a distinct strong enrichment for AS events that generate alternative protein isoforms, particularly among tissue-specific non-IR events. These observations are in line with previous proposals [57, 64, 65] and support the notion that animals have evolved a unique ability to expand their proteomic complexity through AS, specifically for molecular cell and tissue type diversification. What are the biological bases underlying these lineage-specific differences? A widely established hypothesis poses that these differences are due to the distinct lifestyles of plants and animals. Plants are sessile organisms and thus need fast and efficient molecular systems to respond *in situ* to changing environmental conditions and stresses. On the other hand, animals can move away from external perturbations, but for this they require highly specialized cellular and anatomical structures, such as those of muscular and nervous systems, which are particularly enriched for tissue-specific isoforms [25]. Thus, each lineage has evolved a unique specialization of the molecular regulatory tools offered by AS in line with its specific development and physiological needs.

## Conclusions

This study consolidates and extends previous observations as well as unveils unexpected novel patterns. First, we confirm that IR is the most common AS event in *A. thaliana*; however, we find non-IR events to be overrepresented across most regulated AS sets, suggesting the function relevance of these events have been likely underrated. Second, we report a potential new mechanism for plant transcriptome adaptation to stress: by extending the 5’ UTR sequences, AS likely modulates uORF production in response to abiotic and biotic stresses. Third, we revealed an unappreciated genomic regulatory hardwiring, by which sets of genes controlled by either transcriptional or post-transcriptional mechanisms exhibit genomic architectural features that likely determine their regulatory mode in a largely mutually exclusive manner. Forth, we provide evidence that a reduction in core spliceosomal activity is likely behind transcriptomic-wide AS remodeling upon abiotic stress. Finally, compared to animals, we show that *A. thaliana* disproportionally uses AS for stress responses in contrast to tissue-specific transcriptomic and proteomic diversification.

## Methods

### Sample collection and grouping for *A. thaliana*

RNA sequencing (RNA-seq) data was downloaded from the NCBI Short Read Archive (SRA). For this purpose, all available 1,491 RNA-seq experiments from *A. thaliana* at Gene Expression Omnibus (GEO) until June of 2019 were browsed. Experiments with read length equal or larger than 50 nucleotides were shortlisted based on biological interest, trying to cover as many tissues and conditions as possible. A quality control step was done to select the final set of samples based on sequencing depth (key for proper AS quantification), percentage of uniquely mapping reads to the genome and transcriptome, and 3′ sequencing bias as estimated by *vast-tools align* (for details, see https://github.com/vastgroup/vast-tools). Samples that performed poorly on any of these features were discarded unless the experiment was unique and considered to have a high biological relevance. These quality indicators are provided in Additional file 1: Table S1, and low quality values are marked in red. Moreover, when available, the expression of molecular markers was checked to confirm the validity of the experiments, especially in those cases in which the quality of the RNA-seq samples were poorer based on the abovementioned features. Moreover, to prevent overrepresentation of IR events, we discarded non-polyA-selected samples unless no other options were available. In total, *PastDB* provides AS and GE quantifications for 516 RNA-seq independent samples from 63 individual studies (Additional file 1: Table S1). To increase read coverage at splice junctions to afford more accurate quantification of AS, biological replicates from the same sample as well as similar samples from the same lab were pooled together after GE and AS quantification using *vast-tools merge* [25]. The validity of these groupings was confirmed using unsupervised clustering of all samples based on GE data. A similar approach was taken to select and process a set of RNA-seq samples for *C. elegans, D. melanogaster* and *H. sapiens* covering abiotic and biotic stress experiments and different adult tissues (Additional file 1: Table S8).

### Quantification of splicing and gene expression levels

We employed *vast-tools* v2.2.2 to quantify AS and steady-state mRNA levels (referred to as GE) from RNA-seq for *A. thaliana* and the three animal species using the following VASTDB libraries: *A. thaliana* (*araTha10*, based on Ensembl Plants v31; vastdb.ath.20.12.19), *C. elegans* (*ce11*, based on Ensembl v87; vastdb.cel.20.12.19), *D. melanogaster* (*dm6*, based on Ensembl Metazoa v26; vastdb.dme.20.12.19), and *H. sapiens* (*hg38*, based on Ensembl v88; vastdb.hs2.20.12.19). GE was quantified using the cRPKM metric (<u>c</u>orrected-for-mappability <u>R</u>eads <u>P</u>er <u>K</u>bp of mappable sequence per <u>M</u>illion mapped reads)[66]. To make the mapping of the different samples more homogeneous, *vast-tools align* maps only the first 50 nucleotides of the forward read mate (if the sequencing is paired-end) to a reference transcriptome. This transcriptome corresponds to a representative annotated transcript per gene; the list of transcripts is provided in Additional file 12: Table S11. cRPKM values were further normalized using a quantile normalization with the *normalizeBetweenArrays* function from *limma* [67]. For AS, *vast-tools* quantifies exon skipping (ES), intron retention (IR), and alternative donor (ALTD) and acceptor (ALTA) site choices. For all event types, *vast-tools* estimates the percent of inclusion of the alternative sequence (PSI) in a given RNA-seq sample using only exon-exon (or exon-intron for IR) junction reads [25, 26]. Moreover, it provides a “quality score” associated with each PSI value, including information about the read coverage supporting the PSI quantification. Throughout the study, and unless otherwise specified, we used a minimum coverage of VLOW (for further details see https://github.com/vastgroup/vast-tools/blob/master/README.md). Furthermore, in the case of IR events, the quality score includes the *p*-value of a binomial test for read imbalance between the two exon-intron junctions (for details, see [28]). Only introns with a non-significant imbalance (*P* > 0.05) were considered for analyses (--p_IR filter in *vast-tools*).

### Identification of PanAS events and quantification of AS genes

To identify PanAS events, i.e. AS events that generate multiple isoforms across most sample types, we followed a similar approach as previously described [25]. First, we ensured a broad expression level of the gene across tissues by requiring AS events to have sufficient read coverage in at least 20 samples out of the 90 samples from the development and tissues sets (“VLOW” coverage score or higher, and with a non-significant imbalance for IR events). From the events that passed this filtering step, we then defined the PanAS events as those with a PSI between 10 and 90 in >80% of samples with sufficient read coverage (code available in https://github.com/vastdb-pastdb/pastdb).

To quantify the percentage of AS, we implemented several approaches. For Fig. 1 and Additional file 2: Figure S2, AS events and alternatively spliced genes were calculated from 169 samples, which correspond to all samples provided in *PastDB* with the exception of the experiments from RNA-processing mutants. Alternatively spliced events were defined as those with either (i) 10 ≤ PSI ≤ 90 in at least 10% of the samples with sufficient read coverage (“VLOW” coverage score or higher, and with a non-significant imbalance for IR events), or (ii) a PSI range (difference between the maximum and the minimum PSI value across samples) of at least 25. For an event to be considered in this analysis, it had to have sufficient read coverage in at least three samples. We then considered as alternatively spliced those genes with at least one AS event fulfilling these criteria. The total number of genes used to calculate the percentage of AS genes corresponded to the number of multi-exonic genes with at least one event of any type that had sufficient read coverage in at least three samples (whether they were considered alternatively spliced or not). For Additional file 2: Figure S2, we performed a saturation analyses by quantifying the percent of AS genes for increasingly larger random subsets of the 169-sample set. The plotted values for each number of samples corresponded to the median and interquartile range of 100 random subsets of samples of that size (iterations). For the comparison between *A. thaliana* and the three animal species (Additional file 2: Figure S24), we selected, for each species, the samples used for the tissue-specific analysis and performed a saturation analyses by randomly probing subsets of samples (from 5 to 30). For each iteration, we defined AS genes as those with at least one event (from all types or for each individual type tested) with a PSI of either 10 ≤ PSI ≤ 90, 20 ≤ PSI ≤ 80 or 30 ≤ PSI ≤ 70. A valid event had to have sufficient read coverage in at least five samples. As above, the total number of genes used to calculate the percentage corresponded to the number of multi-exonic genes with at least one AS event of any type that had sufficient read coverage in at least five samples. The scripts used for these analyses are available in https://github.com/vastdb-pastdb/pastdb.

### Definition of the core sets of regulated AS events

To identify AS events regulated by abiotic or biotic stress (Additional file 4: Table S3), we first selected five experiments for each type of stress comprising a specific stress condition and a matched control (Additional file 3: Table S2). Based on these data, an AS event was included in the abiotic or biotic stress AS core set if (Additional file 2: Figure S5a):

i. the AS event had sufficient read coverage in at least two out of five experiments of the same stress type (i.e. VLOW or higher in at least two pairs of samples, and with a non-significant imbalance for IR events). Moreover, to ensure the selection of ALTA and ALTD events on exons with a minimum inclusion, we required a minimum PSI of the host exon of at least 25 in each analyzed sample using the option --min_ALT_use 25 of *vast-tools compare*; and
ii. the inclusion level of the AS event was regulated (|ΔPSI| > 15) in the same direction between stress and control conditions for at least two out of five experiments of the same stress type. Events regulated in more than one experiment of the same stress type but in opposite directions were considered ambiguous and not included in the AS core set.

Similarly, to identify tissue-specific AS events (Additional file 4: Table S3), we followed a similar procedure as the one described in [27], in which each tissue is compared to all others. We defined an event to be specifically included or skipped in a given tissue type (Additional file 2: Figure S5b) if:

i. the AS event had sufficient read coverage (‘VLOW’ coverage score or higher, and with a non-significant imbalance for IR events) in at least two replicates of that tissue type and in at least two replicates of at least three other tissue types (Additional file 3: Table S2); and
ii. the average PSI of the AS event in that tissue type is at least 15% higher or lower (i.e. |minimum ΔPSI|>15) than in any other unrelated tissue and the difference in PSI between the target tissue and the average of the other tissues has to be of at least 25 (i.e. |global ΔPSI|>25).

Therefore, although the stress and tissue cores were obtained using comparable approaches, it should be noted that their definition cannot be identical (Additional file 2: Figure S5): whereas for the stress cores we compared pairs of experiments (a stress vs. a control sample), in the case of tissues we compared each tissue against all others for which an event had sufficient read coverage. Further details and code for these analyses are available in https://github.com/vastdb-pastdb/pastdb.

### Definition of the core sets of regulated genes at the GE level

To define the three core sets of regulated genes at the GE level, we used the same samples as per the AS core sets (Additional file 3: Table S2). Then, we implemented a comparable approach to that used for AS (Additional file 2: Figure S5). For the abiotic and biotic stress core sets, we required that a gene changed in expression (FC > 2) in the same direction between stress and control conditions for at least two out of five experiments of the same stress type. Genes that change in opposite directions in experiments of the same stress type were considered ambiguous and not included in the GE core set. Only genes with an expression level of cRPKM ≥ 5 and at least 50 mapped reads in at least two samples from at least two different experiments of the same stress type were considered. To identify tissue-specific genes, we first calculated the median expression levels for each tissue type. Then, for those genes with a minimum median expression of cRPKM ≥ 5 in at least one tissue type, we defined them as tissue-specific if they had a fold change of at least 3 in the same direction respect to all other tissue types and a fold change respect to the median of the other tissues of at least 5. Using TPMs instead of cRPKMs (obtained using the option --TPM in *vast-tools combine*) yielded very similar gene expression core sets (99% overlap for abiotic stress, 98% for biotic stress and 90% for tissues).

### Definition of control groups for each core set

For comparison, we defined two main groups of AS events: a background set (“Genome”) and a non-regulated subset (“AS-NR”). First, we defined the Genome group for each AS core set as the events that passed the same coverage criteria and filters described above to define each core set (see above) but without any requirement regarding PSI values. Second, we obtained an AS-NR group from these background sets for each AS core set. For the stress core sets, AS-NR events were those with a 10 ≤ PSI ≤ 90 in at least one sample (e.g. control or stress) from any of the five experiments of the stress type, and a |ΔPSI| < 5 in all experimental comparisons of the same stress type for which both control and stress samples passed the coverage filters. For the tissue-specific AS core set, AS-NR events were those that were alternatively spliced (10 ≤ average PSI ≤ 90) in at least two tissue types, and that showed a |ΔPSI| < 5 for each tissue type versus the rest. Then, to obtain a common set of Genome and AS-NR events, we selected those AS events that were part of the Genome or AS-NR groups, respectively, for two out of the three AS core sets. In addition, if an AS event was in the AS-NR group of two AS core sets, but it was part of the third AS core set, it was also discarded. In the case of the GE core sets, we also similarly defined a common Genome set of genes by selecting those genes that passed the minimal expression and coverage requirements for at least two of the three comparisons (abiotic stress, biotic stress and tissue specificity).

### Predicted protein impact and sequence feature analysis

Impact of inclusion/exclusion of the alternative sequence for each event was predicted largely as previously described [26], by taking into account the position of the sequence respect to the coding sequence (CDS), the information from annotated isoforms, and whether or not it introduces an in-frame stop codon or a frame shift when included or excluded. Six major groups were defined for all AS events, in which the alternative sequence: (i) is in the 5′ UTR, (ii) is in the 3′ UTR, (iii) generates alternative protein isoforms upon inclusion/exclusion, (iv) disrupts the ORF upon sequence inclusion, (v) disrupts the ORF upon sequence exclusion, and (vi) impacts the CDS with uncertain impact. The latter category corresponds to a minority of cases associated with complex events (including annotated alternative promoters or polyadenylation sites) and were excluded from the analyses. For ES events, in addition to the conditions described in [26], an event that affected the last exon-exon junctions and it was not predicted to elicit NMD (based on the >50-nt rule) was also considered to disrupt the ORF if it was predicted to generate a protein with a truncation of >20% of the length of the reference protein or >300 aminoacids. ORF predictions for all AS events can be retrieved in the Downloads section of *PastDB* (v3; http://pastdb.crg.eu/wiki/Downloads).

To compare exon and intron features associated with splicing regulation, we used *Matt* v1.3.0 [45]. For each compared group of ES and IR events, *Matt cmpr_exons* and *cmpr_introns* automatically extracts and compares multiple genomic features associated with AS regulation, including exon and intron length and GC content, splice site strength, branch point number, strength and distance to the 3′ splice site using different predictions, length and position of the polypyrimidine tract, etc. For the calculation of splice site and branch point strength, we used the available human models. As a complementary approach to the human models, we built position-weighted matrices (PWM) from the alignments of all 3′ (20 nt from the intron and 3 nt from the exon) and 5′ (3 nt from the exon and 6 nt from the intron) splice sites annotated in Ensembl Plants v31 employing the consensus matrix function in the *Biostrings* R library. Then, for each splice site, we used the similarity to these PWMs as an estimate of their strength [4] (code available on https://github.com/vastdb-pastdb/pastdb). Using these values instead of the human-based *MaxEntScan* scores yielded very similar results (Additional file 2: Figure S32). For ALTA and ALTD events, we run *Matt cmpr_exons* providing the exon coordinates with the specific ALTA or ALTD variants as test exons. Finally, for extracting exon-intron structure statistics (including for Additional file 2: Figure S23), we used the following Ensembl annotations: *A. thaliana*, TAIR10 from Ensembl Plants v31; *C. elegans*, WBcel235 Ensembl v87; *D. melanogaster*, BDGP6.22 Ensembl Metazoa v26; and *H. sapiens*, hg38 Ensembl v88.

### Relative contribution of tissues and stress conditions to global PSI variation

For the comparison of the relative contribution to the total PSI variation of the tissue samples versus those of the abiotic or biotic stress experiments, we first calculated for each AS event: (i) the PSI range across tissue types (i.e. the difference between the median PSI of the tissue types with the highest and lowest values), (ii) the highest absolute ΔPSI among the abiotic stress experiments, and (iii) the highest absolute ΔPSI among the biotic stress experiments. The sum of these three values was considered the global PSI variation and the percent of the contribution of each type of variation (tissues, abiotic stress and biotic stress) was calculated. Then, we filtered out the AS events that did not have sufficient read coverage for at least 4 tissue types, 4 abiotic experiments (i.e. both control and stress condition) and 4 biotic experiments and whose global PSI variation was lower than 10. Further details and code for this analysis are available in https://github.com/vastdb-pastdb/pastdb.

### Gene Ontology enrichment analyses

To identify significantly enriched biological processes, molecular functions and cellular components analyses among different sets of genes, we used Gene Ontology (GO) enrichment analyses as implemented by the functional annotation classification system DAVID version 6.8 [68]. For each comparison, only genes with events that passed equivalent read coverage filters as those being tested were used as a background.

### Evolutionary analyses of AS core sets

To analyze the level of genome conservation [39, 69] for each AS core set and event type within *Brassicaceae*, we first generated chain genome alignments (liftOver files) between *A. thaliana* and four closely related species: *Arabidopsis lyrata* (v.1.0, estimated split 7.1 million years ago [MYA] [70]), *Camelina sativa* (Cs, 9.4 MYA), *Arabis alpina* (A_alpina_V4, 25.6 MYA) and *Brassica rapa* (Brapa_1.0, 25.6 MYA). For this purpose, we downloaded the genome sequences for these four species from Ensembl Plants v48, and generated the chain alignments following the UCSC pipeline and using *blat* with - minIdentity=80 and -minScore=50 parameters (the full pipeline is available on https://github.com/vastdb-pastdb/pastdb). Next, we lifted the coordinates for all *vast-tools* events from *A. thaliana* by AS type and assessed their presence in each of other four genomes (“genome conservation”). Following previous studies [25], for alternative exons, we further required that the lifted exonic sequence was surrounded by at least one canonical 5′ (GT/C) or 3′ (AG) splice site dinucleotide in the target species. For alternative 5′ and 3′ splice site choices we required the canonical splice site dinucleotide to be conserved. Finally, for IR events, we required the lifted region to intersect with an annotated intron (from the Ensembl Plants v48 annotations) in the target species, using *bedtools intersect*. These results are shown in Additional file 2: Figure S10.

To assess the overlap among the AS core sets from *A. thaliana* and the three animal species, we used *ExOrthist* v0.0.1.beta (https://github.com/biocorecrg/ExOrthist). This software is designed to identify orthologous exons based on intron position-aware pairwise protein alignments (as previously described in [27, 31]). *ExOrthist* uses clusters of gene orthologs, which were obtained using *Broccoli* [71] with the sets of translated reference transcript per gene for *A. thaliana, C. elegans, D. melanogaster*, and *H. sapiens*. Clusters of orthologous exons were then obtained for all annotated exons plus all *vast-tools* exons (added through the --extraexons argument) for each species using the default settings for ‘long evolutionary distance’ for all pairwise species comparisons (*long_dist*, minimum exonic sequence similarity [ex_sim] = 0.20, maximum ratio difference between exon lengths [ex_len] = 0.65 and overall protein sequence similarity [prot_sim] = 0.15; see https://github.com/biocorecrg/ExOrthist for further details). Finally, to generate the four-way Venn diagrams in Additional file 2: Figure S29, we assessed the overlap of each core set of AS events from each species with the others at two levels (code available in https://github.com/vastdb-pastdb/pastdb). First, we checked if any orthologous gene (defined as being in the same gene cluster) in the target species also hosted an AS event (of any type) belonging to the same core set (i.e. abiotic stress, biotic stress or tissues). Second, we assessed whether the exact splice site(s) affected by the AS event were also affected by an AS event (of any type) of the same core set in the target species. In the case of 5′ UTR AS events, these were conservatively considered to fall in the same splice site if orthologous genes in the two species had a core set AS event in the 5′ UTR. It should be kept in mind that these overlaps do not imply evolutionary conservation; rather, for such distantly related species, the overlaps are likely the result of convergent evolution.

### *PastDB* website, database and data sources

The website was built over a LAMP environment, where MariaDB is used as DataBase Management System for *PastDB*, while MediaWiki works as Content Management System platform. MediaWiki BioDB extension, a custom modification of a pre-existing extension named External Data, was installed for reusing data directly from MySQL tables. To allow faster custom searches (by gene name or ID, Event ID or coordinate) selected search fields were mapped into document structures, which were imported into an IBM CouchDB 3 (a NoSQL document DBMS) server instance. Predefined indexes were created for gene descriptions (taking advantage of IBM CouchDB 3 Lucene capabilities) and for coordinates (including chromosome, start, end and strand fields). After that, a custom MediaWiki extension previously created for *VastDB* [25], and some JavaScript functions were created in order to enable forms and other specific needs of the website. Coordinate search results were also coupled to an embedded genome browser view.

For the embedded genome browser *PastDB* uses a custom UCSC browser instance (http://ucsc.crg.eu). This browser includes a general transcript track (all transcripts annotated in the GTF based on TAIR10 obtained from Ensembl Plants v31), a gene track (with the single reference transcript per gene; Additional file 12: Table S11) and the *PastDB* AS events track. The latter shows, using a color scheme based on the AS type, the AS events with the highest PSI variation across all 202 *PastDB* samples. Nonetheless, *PastDB* includes all the information for all AS events that are quantified by *vast-tools*. For comparison, we matched *atRTD2* AS annotations [14] with those provided in *PastDB*. The four major types of AS events were extracted from *atRTD2* using *SUPPA2* [72] and compared to *PastDB* events using splice site coordinate match. Despite substantial differences in the methods and RNA-seq samples used to build each resource, 84.4% of exons, 75.3% of introns, 82.4% of acceptor sites and 72.0% of donors annotated as alternatively spliced in *atRTD2* are present in *PastDB*; conversely, 59.2%, 37.5%, 65.4% and 54.5% of highly alternatively spliced events of each respective type across the main *PastDB* samples are annotated as alternatively spliced in the *atRTD2* genome annotation.

Gene expression and PSI data were represented using Rickshaw, a D3.js-based JavaScript library [http://code.shutterstock.com/rickshaw/]. By using this JavaScript library instead of pre-computed plots, users can choose which data they are shown, from a range of read stringency thresholds and biological samples. In the case of “Special datasets”, sets of samples related to abiotic stress, biotic stress, light conditions, and RNA-processing factors, the values are shown as static PNGs that can be downloaded. Finally, to calculate the overlap with protein domains and intrinsically disordered regions, we used PfamScan (version 2.3.4) [73] and Disopred3 [74], respectively. Plots for domain overlap of one inclusion and one skipping isoform were done using custom R scripts and are static PNG images automatically uploaded when available. All datasets included in *PastDB* can be downloaded from http://pastdb.crg.eu/wiki/Downloads.

## Abbreviations

AS: alternative splicing;
GE: gene expression;
ALTD: alternative splice donors;
ALTA: alternative splice acceptors;
ES: exon skipping;
IR: intron retention;
CDS: coding sequence;
UTR: untranslated region;
ORF: open reading frame;
NMD: nonsense-mediated RNA decay.

## Authors’ contributions

GM and MI conceived the project and designed the analyses. GM, YM, FM and MI performed computational analyses. GM, YM, PD and MI contributed to the interpretation of results. GM and MI generated the data for *PastDB*. GM, MI and PD wrote the manuscript. All the authors critically reviewed and approve the final manuscript.

## Acknowledgements

We thank Toni Hermoso from the BioCore at CRG for his work developing and maintaining *PastDB* and the custom UCSC browser, Xavi Grau-Bove and Andre Gohr for their help with specific analyses, and Nuno Barbosa-Morais for useful suggestions.

## Availability of data and materials

All quantification and genome features and annotations used for all analyses can be accessed and downloaded through *PastDB* (pastdb.crg.eu).

