## Supplemental Information for "Alternative splicing landscapes in *Arabidopsis thaliana* across tissues and stress conditions highlight major functional differences with animals"

**<u>Supplementary Materials for:</u>**

Supplementary Tables:

**Table S1.** ***A. thaliana*** **RNA-seq datasets compiled for** ***PastDB*****.** Details of the 516 samples obtained from the NCBI Short Read Archive (SRA). The quantification of mRNA steady-state levels and alternative splicing of these samples are provided in the website *PastDB* (www.pastdb.crg.eu). Information includes i) SRA identifier, ii) original name, iii) *PastDB* groupings, iv) number of reads and length, v) percentage of reads mapping unique and multiple times at the genome or reference transcriptome (Table S11), vi) estimation of 3′ sequencing biases (for details, see https://github.com/vastgroup/vast-tools), and vii) source of the data. Low quality values for uniquely genome mapping (<40%), uniquely transcriptome mapping (<50%), and 3′ sequencing biases (<1) are highlighted in red font.

**Table S2.** ***A. thaliana*** **RNA-seq datasets used to define the tissue-specific and stress-responsive GE and AS core sets.** Details of the 108 samples obtained from the NCBI Short Read Archive (SRA) used to define the tissue-specific and stress-responsive GE and AS core sets in *A. thaliana*. Information includes: i) SRA identifier, ii) original name, iii) PastDB groupings, iv) number of reads and length, and v) source of the data.

**Table S3. List of AS events differentially regulated in response to abiotic and biotic stimuli or in different tissues of** ***A. thaliana*****.** Differentially regulated AS events in each of the five experiments selected to assess biotic or abiotic responses, as well as in any of the six *A. thaliana* tissues tested. AS events belonging to stress and tissue AS core sets are indicated. Information for each event includes: i) Locus, ii) *vast-tools* event identifier, iii) genomic coordinates, iv) type of event, and v) percent of inclusion (PSI) — quantification based on output files generated by *vast-tools*. NA for PSI values indicates insufficient read coverage (N coverage score) or, in the case of IR events, a significant read imbalance between both intron-exon junctions. See *vast-tools* README for further details (https://github.com/vastgroup/vast-tools).

**Table S4. List of genes differentially regulated in response to abiotic and biotic stimuli or in different tissues of** ***A. thaliana*****.** Differentially regulated genes in each of the five experiments selected to assess biotic or abiotic responses as well as in any of the six *A. thaliana* tissues tested. Genes belonging to stress and tissue GE core sets are indicated. For each gene, expression values using the cRPKM metric are provided. cRPKM values were obtained using *vast-tools*. NA for gene expression values indicates that in neither of the two samples of a given experiment (e.g. control and stress) did the gene have a minimal expression of at least 5 cRPKMs and/or a raw read count of at least 50 (see Methods for details).

**Table S5. Enriched GO terms for genes belonging to the abiotic, biotic and tissue AS and GE core sets in *A. thaliana*.** Complete results of each GO enrichment analysis obtained from the functional annotation classification system DAVID [1].

**Table S6. List of samples related to *A. thaliana* RNA-processing factor mutants.** List of samples used in the analysis shown in Figure 4E and Supplementary Figure 20. Samples were obtained from the NCBI Short Read Archive (SRA). For each sample, i) SRA identifier, ii) original name, iii) *PastDB* groupings, iv) number of reads and length, and v) source of the data are provided.

**Table S7. RNA-seq analyses of RNA-processing factor mutant experiments.** The effects of mutations in each studied RNA-processing factor on AS events of each AS core set are summarized, either for each type of AS event or all AS events combined. "CORE": type of core set. "RBP_Exp": RBP mutant investigated. Comparisons of loss-of-function mutants in different conditions with respect to the control, with the exception of *grp7* (overexpression vs. WT) and *sr45* (overexpression vs. mutant), were performed. "N_events (total)": number of AS events with sufficient read coverage in both control and experimental condition. "UP_in_Exp": number of AS events with a $\Delta PSI > 15$ in the mutant. "DOWN_in_Exp": number of AS events with a $\Delta PSI < -15$ in the mutant. "UP-UP", "DOWN-DOWN", "UP-DOWN" and "DOWN-UP": number of AS events that are up- or down-regulated in the specific core set (first) and up- or down-regulated in the mutant (second). "% same direction": percentage of AS events changing in the same direction (i.e. UP-UP and DOWN-DOWN) with respect to the total changing events. "P-val (raw)": raw *p*-value of a Binomial test for the number of events changing in the same direction vs. the total. "P-val (corr)": Bonferroni-corrected p-value by event AS type (i.e. each raw *p*-value was multiplied by 54). "N_events (core_UP/core_DOWN/AS_NR)": number of AS events of each subset that have sufficient read coverage in control and mutant. "% changing (core_UP/core_DOWN/AS-NR)": percentage of AS events that changed in any direction ($|\Delta PSI| > 15$) in the mutant condition.

**Table S8. RNA-seq datasets used to define the tissue-specific and stress-responsive AS event core sets in *C. elegans*, *D. melanogaster* and *H. sapiens*.** Details of the 460 samples obtained from the NCBI Short Read Archive (SRA) used to define stress and tissue core sets in the different animal organisms. Information includes: i) SRA identifier, ii) original name, iii) *VastDB* groupings, iv) number of reads and length, and v) source and information about data.

**Table S9. List of AS events belonging to the stress and tissue AS core sets in *C. elegans*, *D. melanogaster* and *H. sapiens*.** AS events belonging to stress and tissue AS core sets are indicated. Information for each event includes: i) Locus, ii) *vast-tools* event identifier, and iii) genomic coordinates and related information.

**Table S10. Enriched GO terms for genes belonging to the abiotic, biotic and tissue AS core sets in *C. elegans*, *D. melanogaster* and *H. sapiens*.** Complete results of each GO enrichment analysis obtained from the functional annotation classification system DAVID [1].

**Table S11. Representative transcript per gene used for GE quantification in *vast-tools* and other analyses in this study.** Reference transcript per gene from *A. thaliana*; TAIR10 annotation from Ensembl Plants v31.

### Supplementary Figures:

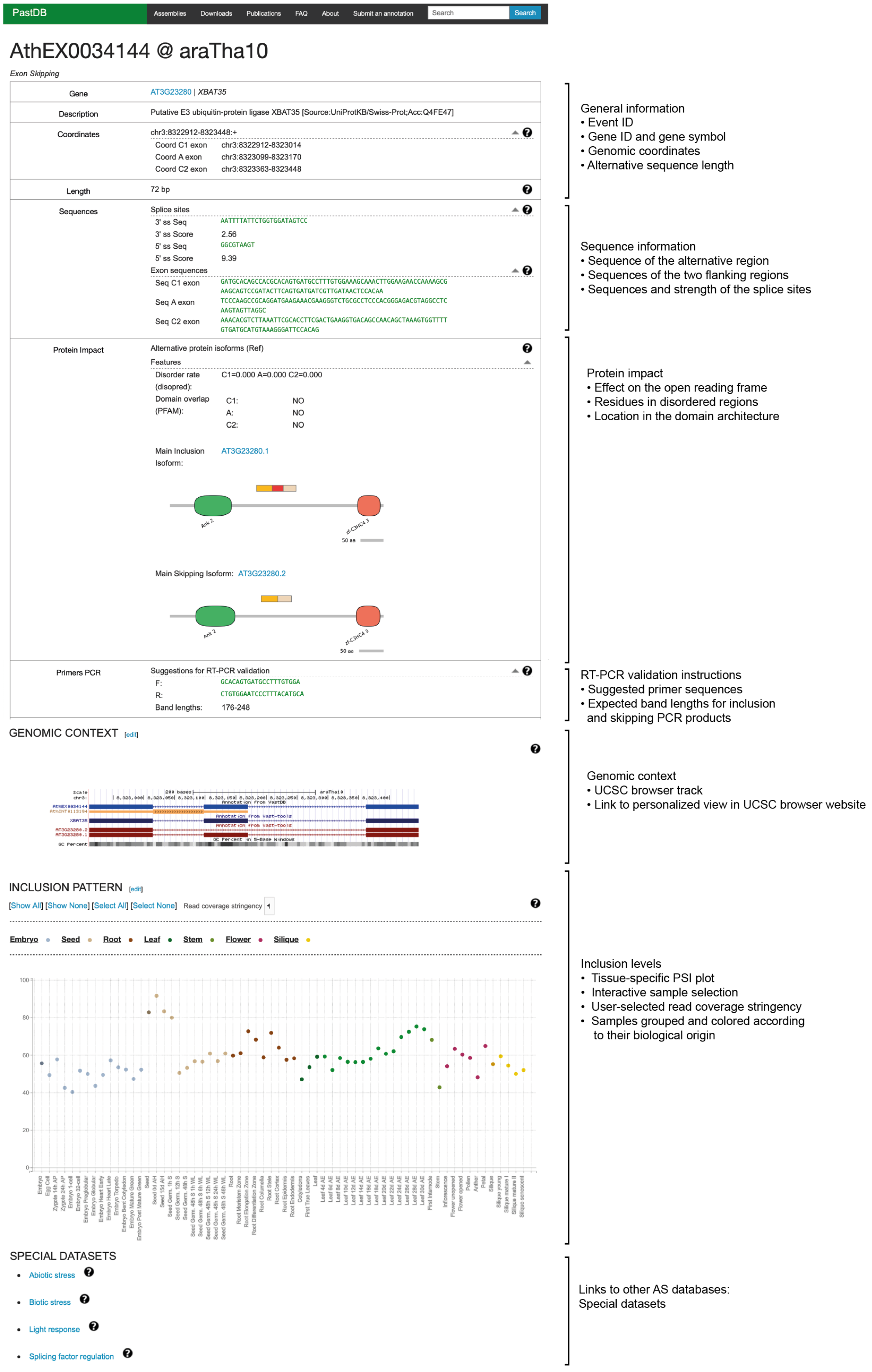

**Figure S1. Example of an AS event view entry from *PastDB***

Adapted screenshot of the Event view from *PastDB* (http://pastdb.crg.eu) for a plant AltEx event (based on AthEX0034144). Event view includes, from top to bottom, general information, sequence information, protein impact of the alternative sequence, suggested primers for RT-PCR validation, genomic context (linking to UCSC Genome Browser), interactive plot of PSI across 64 plant cell and tissue types from averaging 90 samples, plots of PSIs for other datasets of special interest, links to other AS databases, and mapping to protein structures (when available).

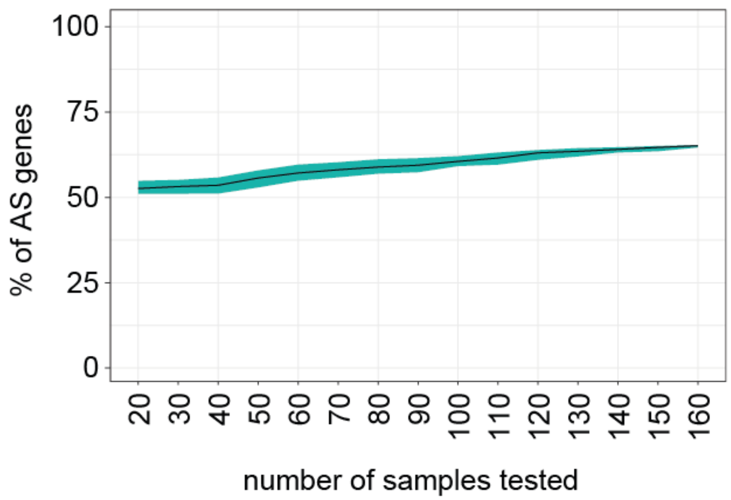


**Figure S2. Quantification of alternatively spliced genes in *A. thaliana* by subsampling RNA-seq datasets.** Percentage of multi-exonic genes harboring at least one AS event with a PSI between 10 and 90 in at least 10% of the tested samples and/or a PSI range ≥ 25. For each number of samples tested, both the median and the interquartile range of 100 iterations selecting random subsets of samples are shown.

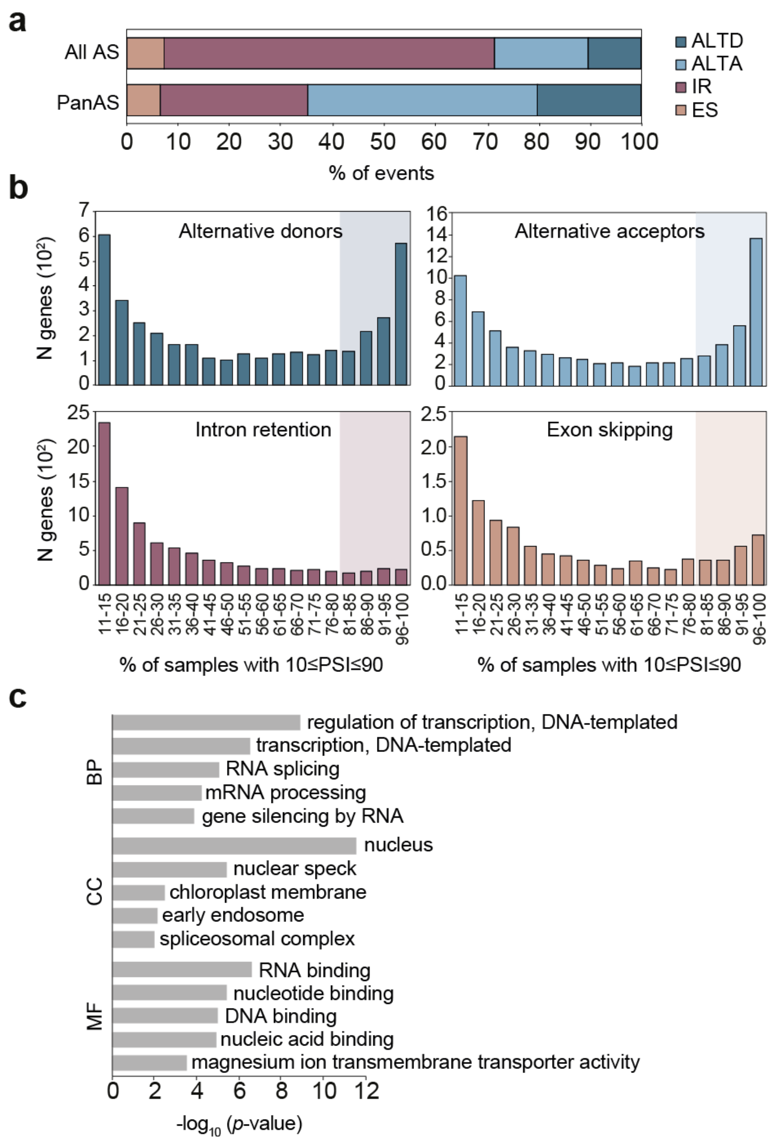


**Figure S3. Identification and characterization of PanAS events in *A. thaliana*. a** Distribution of types of AS among PanAS events in comparison with the genome-wide distribution of AS events shown in Figure 1. **b** Number of each type of AS event based on the percentage of samples in which the AS event has a PSI between 10 and 90. PanAS events correspond to the last four bars: AS events alternatively spliced in more than 80% of the samples. **c** Top five most enriched Gene Ontology (GO) categories for biological process (BP), cellular component (CC) and molecular function (MF) among genes harboring PanAS events. *p*-values (Fisher's Exact test) were obtained from DAVID. ALTD, alternative splice donor; ALTA, alternative splice acceptor; ES, exon skipping; IR, intron retention; N, number.

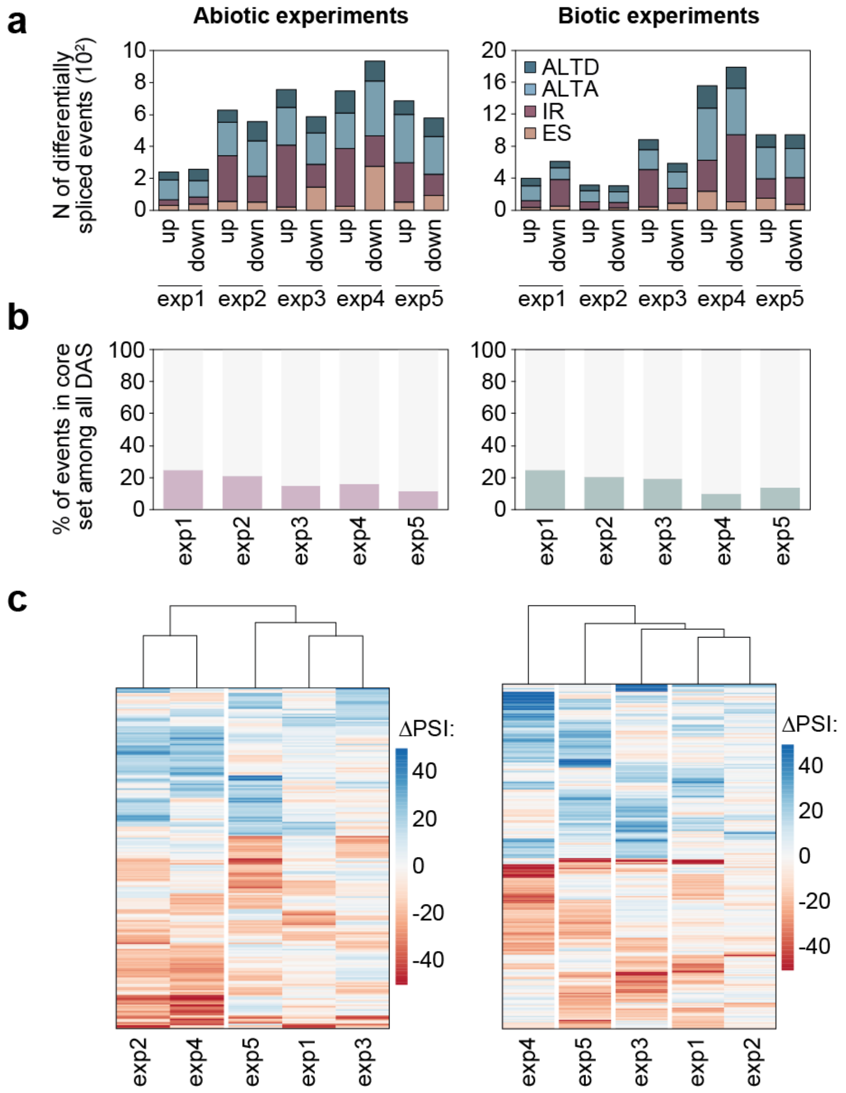


**Figure S4. Definition of a core set of AS events responding to abiotic or biotic stress. a** Number of events by AS type whose alternative sequence is differentially up or down regulated in each abiotic or biotic stress experiment. **b** Percentage of AS events differentially spliced in each experiment that belong to the abiotic (left) or biotic (right) core set of events. **c** Heatmap showing the ΔPSI of each differentially regulated AS event between stress and control conditions in each abiotic (left) and biotic (right) experiment. Only AS events with sufficient read coverage in both control and stress conditions in the five experiments are shown for each type of stress (n = 214, abiotic; n = 281, biotic). ALTD, alternative splice donor; ALTA, alternative splice acceptor; ES, exon skipping; IR, intron retention; DAS: differentially spliced event; N, number.

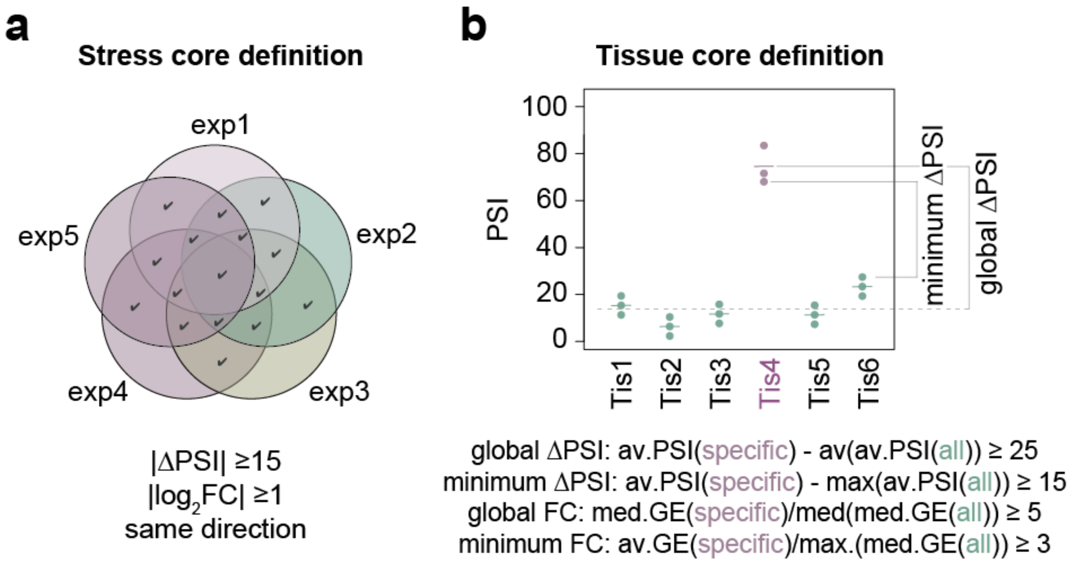


**Figure S5. Schematic representation of the definitions used to identify the core sets of genes and events changing in response to stresses and in tissues. a** Core stress-responding genes/events are defined as those that are differentially expressed/spliced in the same direction in at least two out of five experiments. **b** Tissue-specific genes/events are defined as those that are downregulated or upregulated in one of the six selected tissues. See Methods for details.

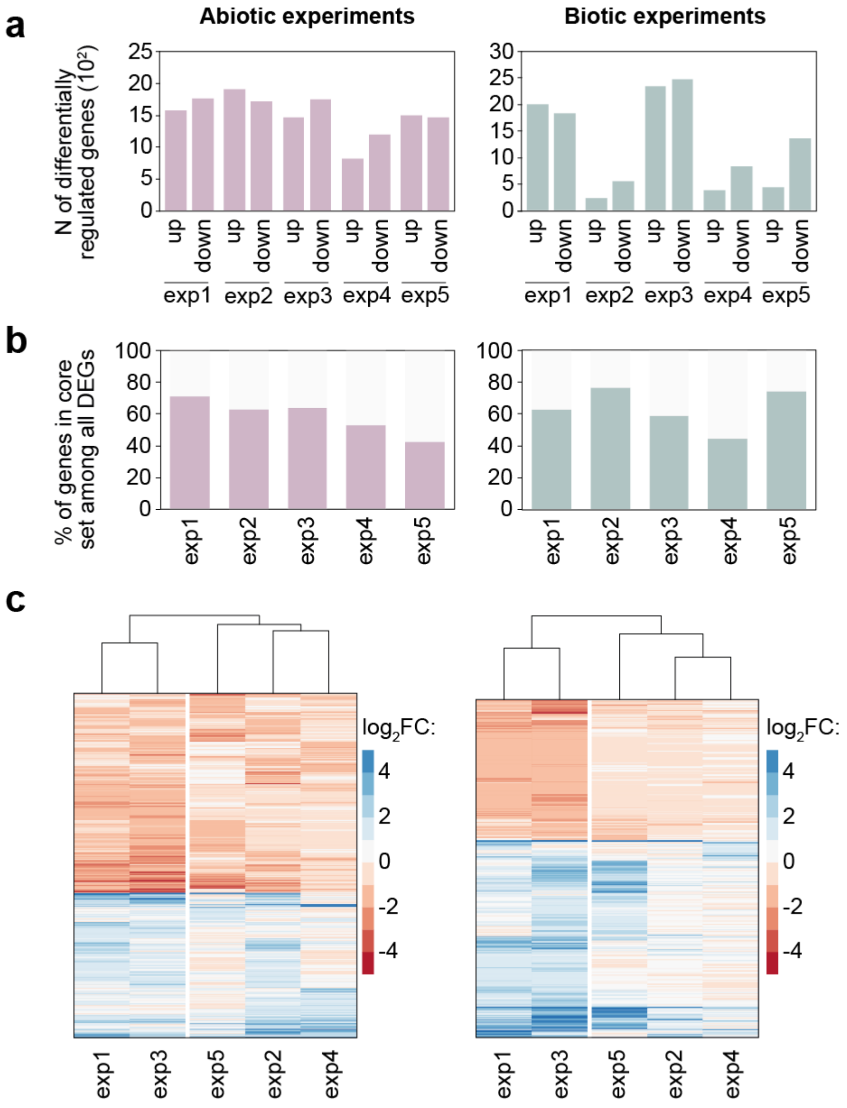


**Figure S6. Core sets of genes with differential GE regulation upon abiotic or biotic stress. a** Number of genes differentially up- or down-regulated in each abiotic or biotic stress experiment. **b** Percentage of differentially expressed genes in each experiment belonging to the abiotic (left) or biotic (right) core sets. **c** Heatmap showing the $\log_2$ fold change (FC) between stress and control conditions in each abiotic (left) and biotic (right) stress experiment. Only genes with a minimum expression (cRPKM≥5 and at least 50 read counts) in control or stress conditions for every abiotic or biotic experiment are depicted (n = 2072, abiotic; n = 1573, biotic). DEG: Differentially expressed gene; N, number.

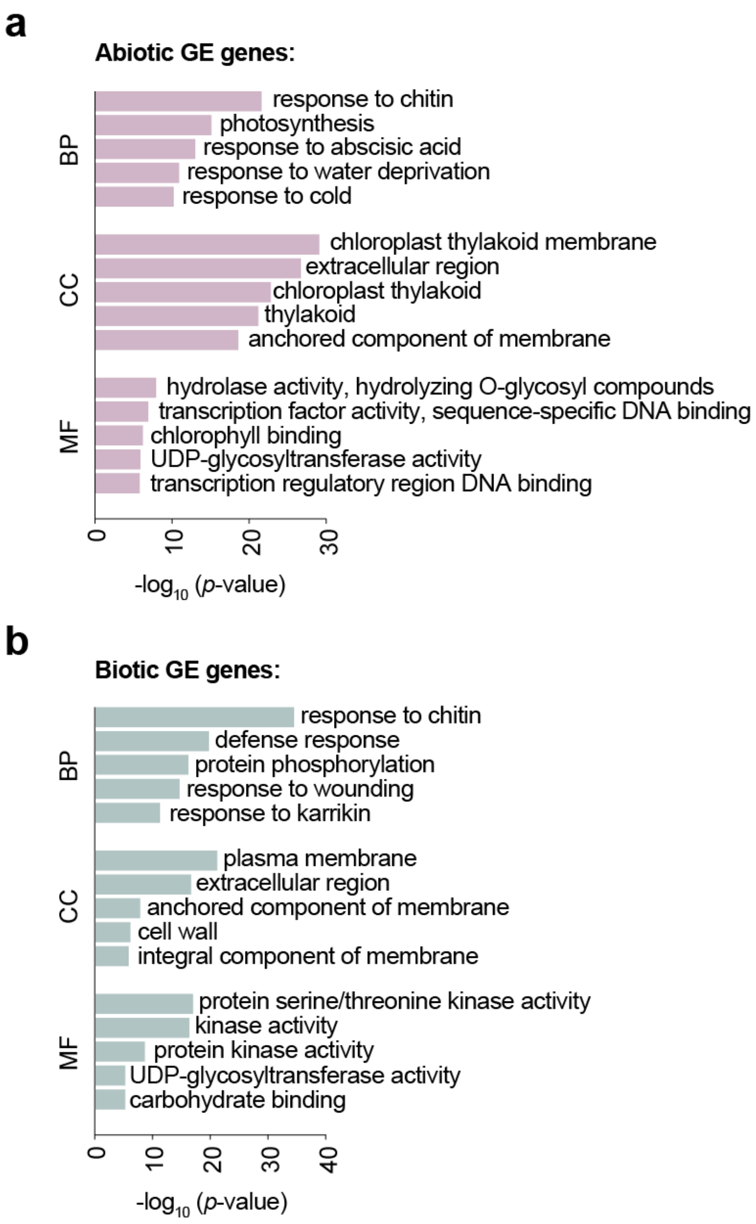


**Figure S7. Functional enrichment analysis for genes belonging to each stress GE core set.** Top five most enriched Gene Ontology (GO) categories for biological process (BP), cellular component (CC) and molecular function (MF) of genes from the abiotic (**a**) and biotic (**b**) expression core sets. *p*-values (Fisher's Exact test) were obtained from DAVID.

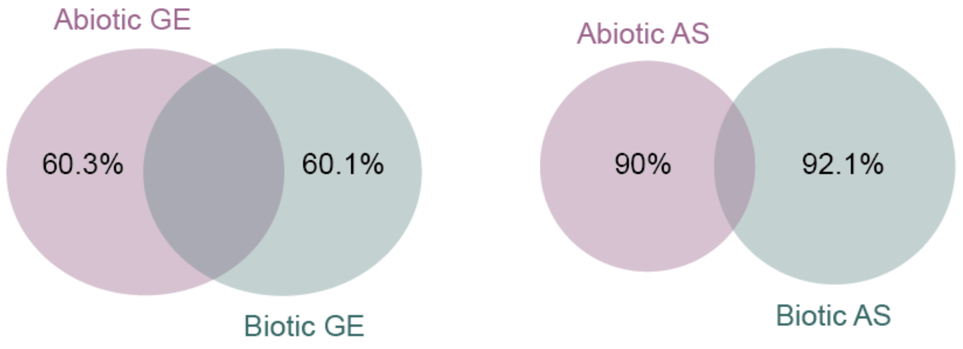


**Figure S8. Overlap between abiotic and biotic core sets.** Venn diagram showing the overlap between the core sets of differentially expressed genes (GE) or spliced (AS) events in response to abiotic (purple) or biotic (green) stimuli.

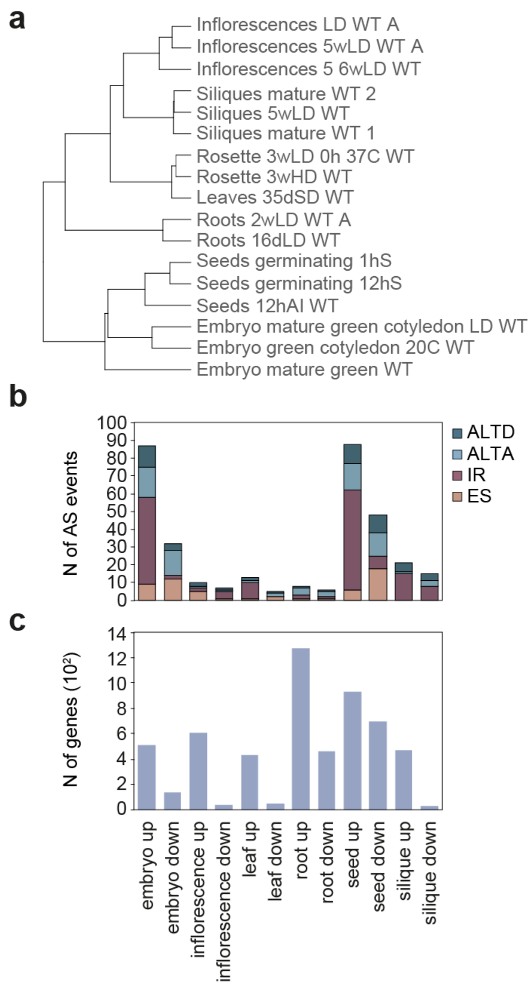


**Figure S9. Tissue-specific AS events and genes. a** Unsupervised clustering of samples used for tissue comparisons based on GE profiles (distance matrix corresponds to 1 - Spearman correlation coefficients). **b, c** Number of AS events sorted by type (**b**) or genes (**c**) that are up- or down-regulated in any of the six studied tissue samples. ALTD, alternative splice donor; ALTA, alternative splice acceptor; ES, exon skipping; IR, intron retention; N, number.

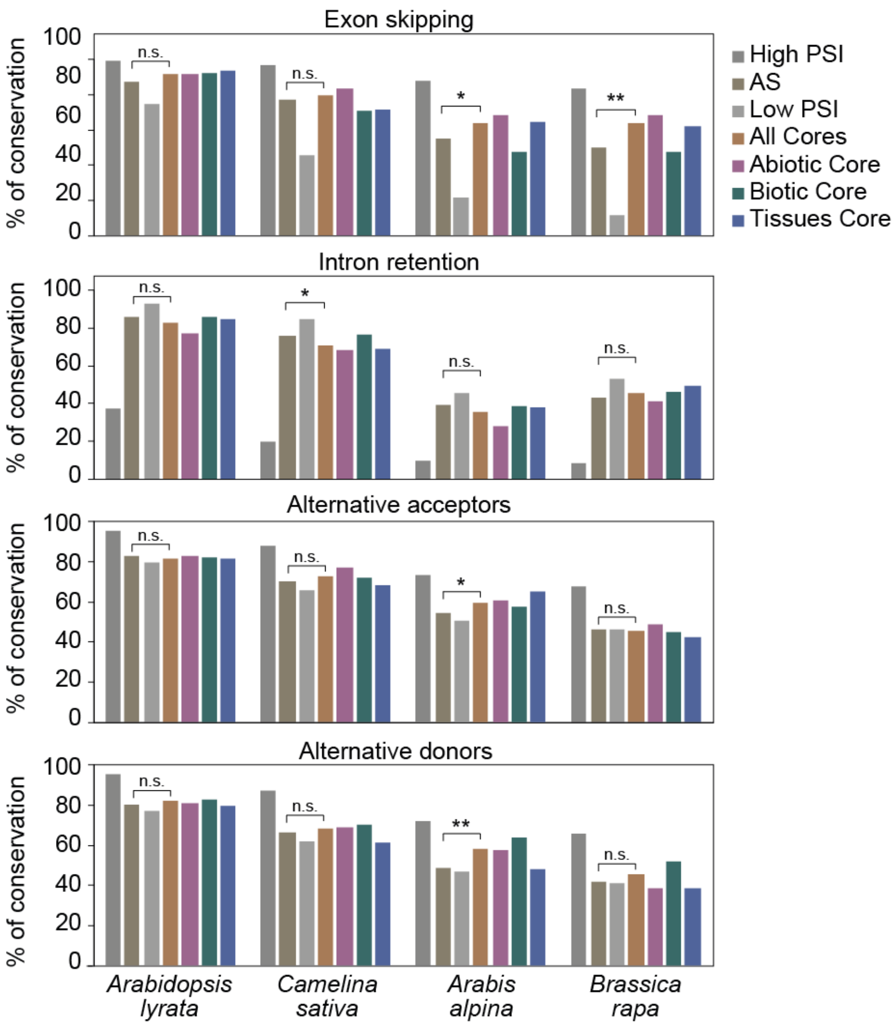


**Figure S10. Genome-conservation of AS core set events by AS type.** Percent of AS events of each type (EX, IR, ALTA and ALTD) from different sets that were found conserved using liftOver in the genomes of *Arabidopsis lyrata* (estimated split 7.1 million years ago [MYA] [2]), *Camelina sativa* (9.4 MYA), *Arabis alpina* (25.6 MYA) and *Brassica rapa* (25.6 MYA). *p*-values indicate statistical significance as assessed by a two-sided Fisher's Exact test comparing the merge of AS core sets and the AS group ($P < 0.05$ (*); $P < 0.01$ (**)). High PSI, events with PSI > 90 across all samples; Low PSI, events with PSI < 10 across all samples; AS, events that are alternatively spliced (10≤PSI≤90 in at least 10% of the samples and/or a PSI range ≥ 25), excluding those in the AS core sets; Core, events in any of the three AS core sets (abiotic, biotic and tissue).

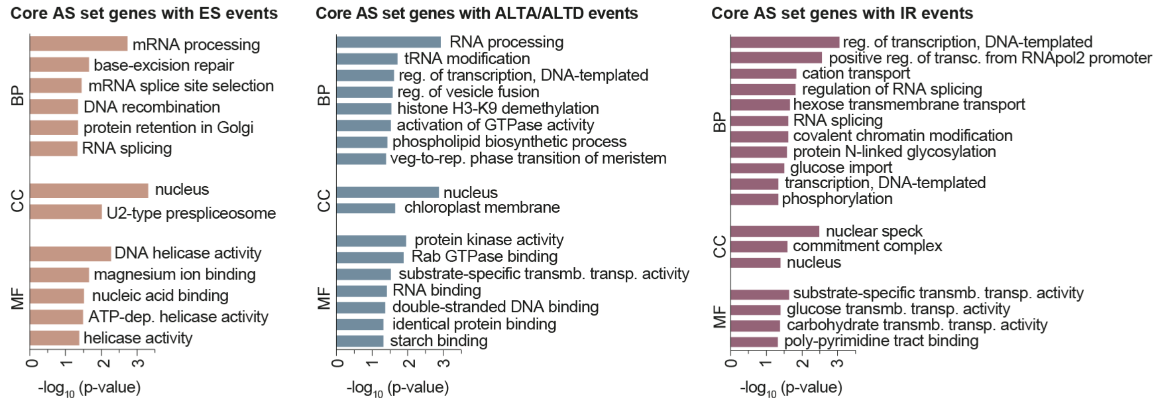


**Figure S11. Functional enrichment analysis of AS-regulated genes in *A. thaliana* classified by type of event.** Enriched Gene Ontology categories of biological process (BP), cellular component (CC) and molecular function (MF) for genes with exon skipping (ES; left), alternative splice sites (ALTD/ALTA; middle) and intron retention (IR; right) events belonging to the three AS core sets. *p*-values (Fisher's Exact test) were obtained from DAVID.

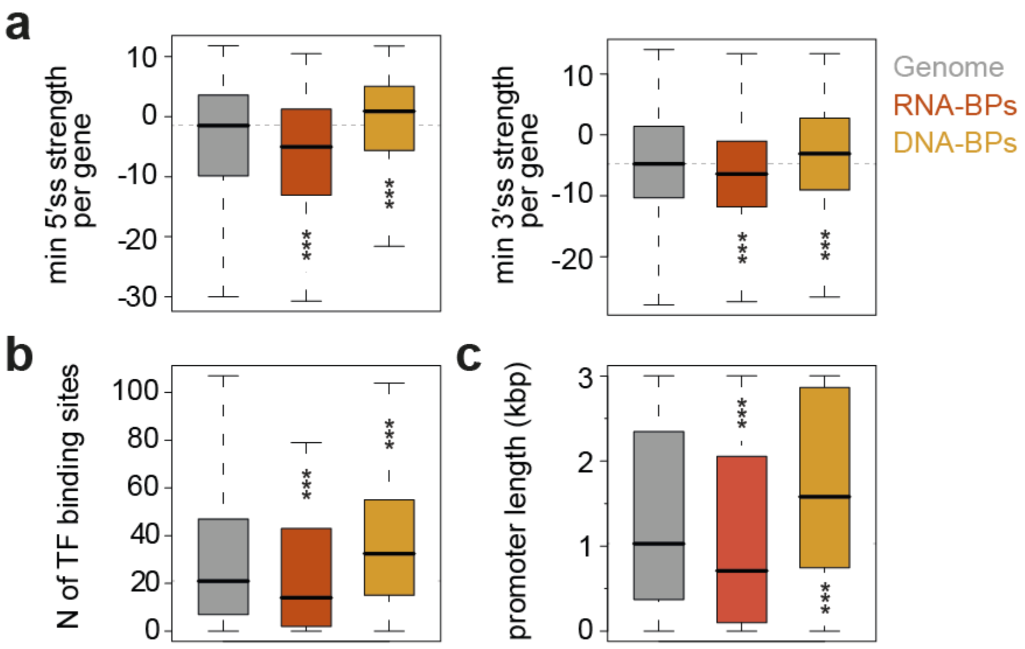


**Figure S12. Genomic features of RNA and DNA binding proteins. a** Distributions of the lowest 5′ splice site (left) or 3′ splice site (right) strength per gene for each category. Values correspond to the splice site with the lowest maximum entropy score in each gene, as calculated in [3] (see Methods). **b** Number of binding sites of *A. thaliana* transcription factors (TF) per gene's promoter. **c** Promoter length for each gene. Promoter information was obtained from [4]. "Genome" shows the distributions of the corresponding values for all multi-exonic *A. thaliana* genes. RNA and DNA binding proteins were downloaded from cisBP-RNA [5] and cisBP [6], respectively. *p*-values indicate significance as assessed by Wilcoxon test ($P < 0.05$ (*); $P < 0.01$ (**); $P < 0.001$ (***)). BPs, binding proteins; N, number.

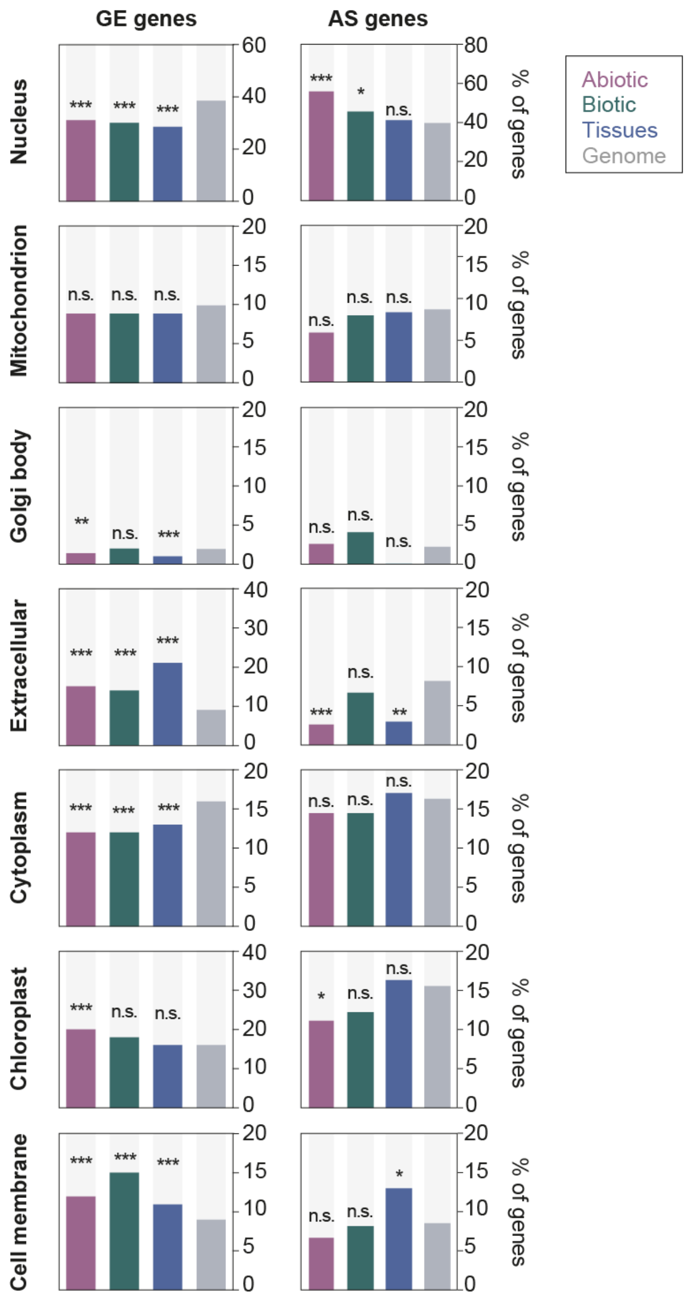


**Figure S13. Subcellular localization of genes belonging to the GE and AS core sets.** Subcellular localization of differentially expressed (GE) or spliced (AS) genes from the abiotic, biotic and tissue core sets based on annotations from TAIR (http://www.arabidopsis.org). *p*-values indicate statistical significance as assessed by two-sided Fisher's Exact tests (P < 0.05 (*); P < 0.01 (**); P < 0.001 (***)). "Genome" contains all *A. thaliana* genes fulfilling the coverage criteria used for the corresponding GE and AS analyses.

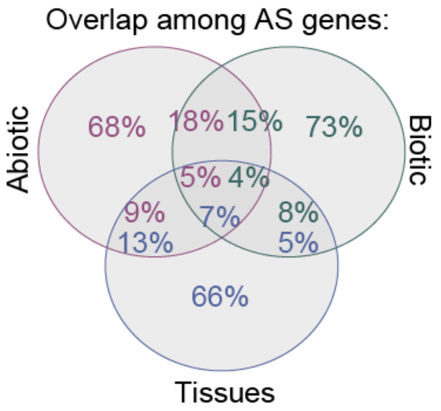


**Figure S14. Overlap among AS genes from each core set.** Venn diagram showing the overlap between differentially spliced genes belonging to the abiotic stress (purple rim), biotic stress (green rim) or tissue-specific (blue rim) core sets.

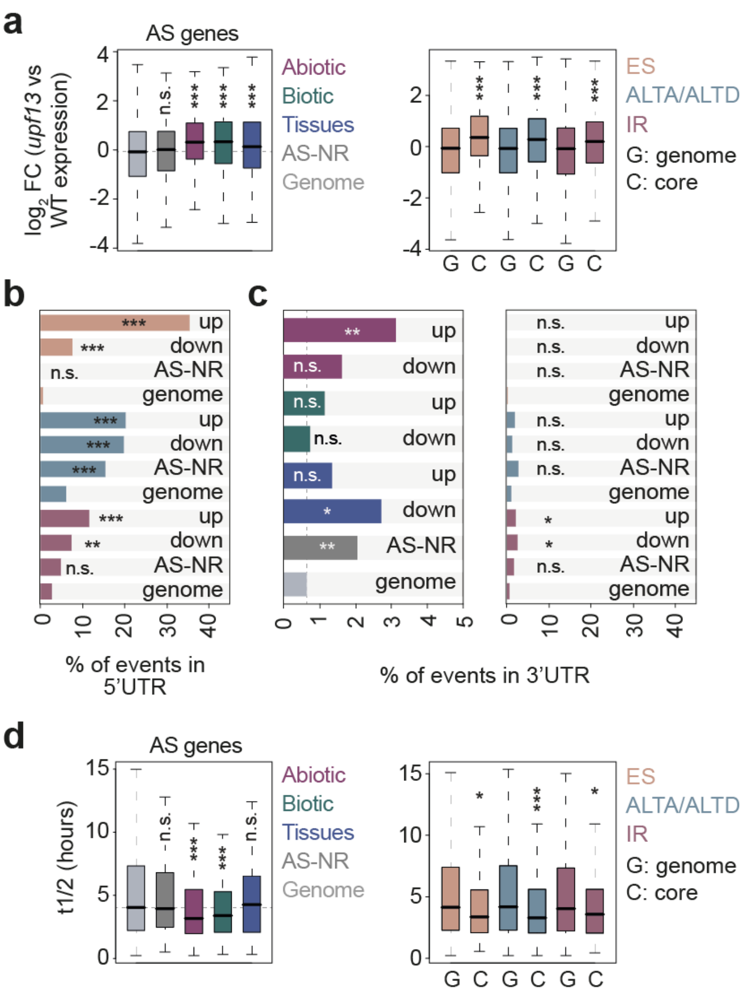


**Figure S15. Impact of regulated AS events on ORF and UTRs. a** $Log_2$ of the fold change in the expression values of *upf13* double mutant compared to WT for different sets of AS genes (left) and by AS type (right). **b** Percentage of events by type of AS that occur in the 5′ UTR of their respective genes. Events from the three AS core sets were grouped according to whether their alternative sequence is up- or down-regulated. **c** Percentage of AS events located in the 3′ UTR of its respective gene by regulated set (left) or by event type (right). **d** mRNA half-life of genes belonging to different sets of AS genes (left) or grouped by AS type (right). *p*-value indicates statistical significance as assessed by Wilcoxon Rank-Sum tests (**a**,**d**) or two-sided Fisher's Exact tests (**b**,**c**) ($P < 0.05$ (*); $P < 0.01$ (**); $P < 0.001$ (***))."Genome" contains all multi-exonic *A. thaliana* genes fulfilling the coverage criteria used for the corresponding AS analysis. Data from **a** and **d** was obtained from [7] and [8], respectively. AS-NR, alternative spliced-non regulated; n.s., not significant.

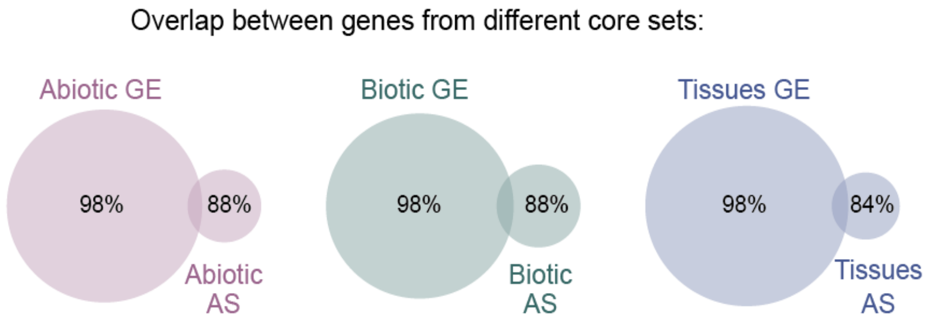


**Figure S16. Overlap between differentially expressed and spliced genes.** Venn diagrams showing the overlap between differentially expressed (GE) and spliced (AS) genes belonging to the abiotic (left), biotic (middle) and tissue (right) core sets.

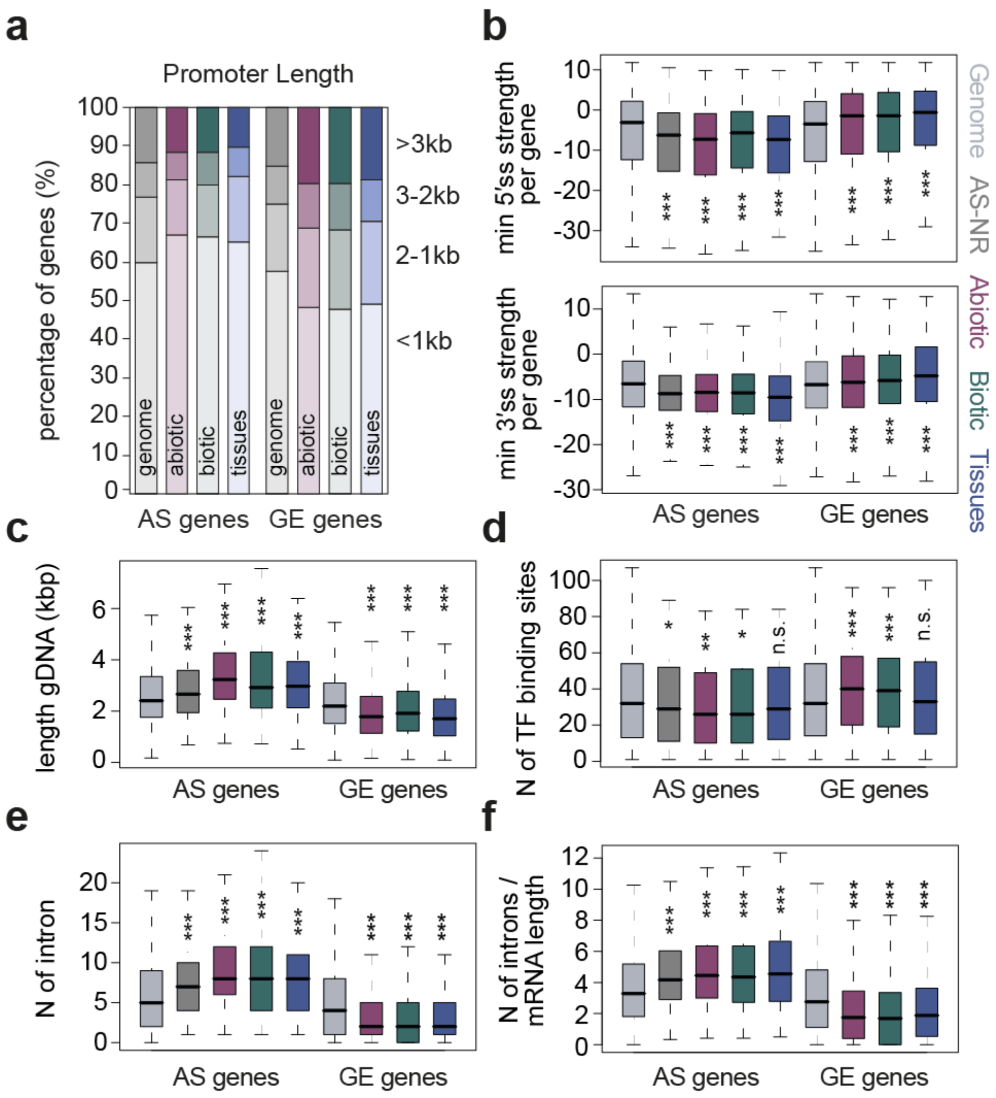


**Figure S17. Genomic features of genes belonging to GE and AS core sets. a** Distribution of genes from the abiotic, biotic and tissue AS or GE core sets according to promoter length. **b** Lowest 5´splice site (top) or 3´splice site (bottom) strength for each gene. **c** Genomic size. **d** Number of binding sites for *A. thaliana* TFs**. e** Number of total introns per gene**. f** Density of introns per kbp of mRNA. Splice site strength was calculated as described in [3] (see Methods). Data from **d** was obtained from https://agris-knowledgebase.org. **a-g** "Genome" contains *A. thaliana* genes fulfilling the coverage criteria used for GE and AS analyses. *p*-values indicate statistical significance as assessed by Wilcoxon Rank-Sum tests ($P < 0.05$ (*); $P < 0.01$ (**); $P < 0.001$ (***)). AS-NR, alternative spliced-non regulated; n.s., not significant; N, number.

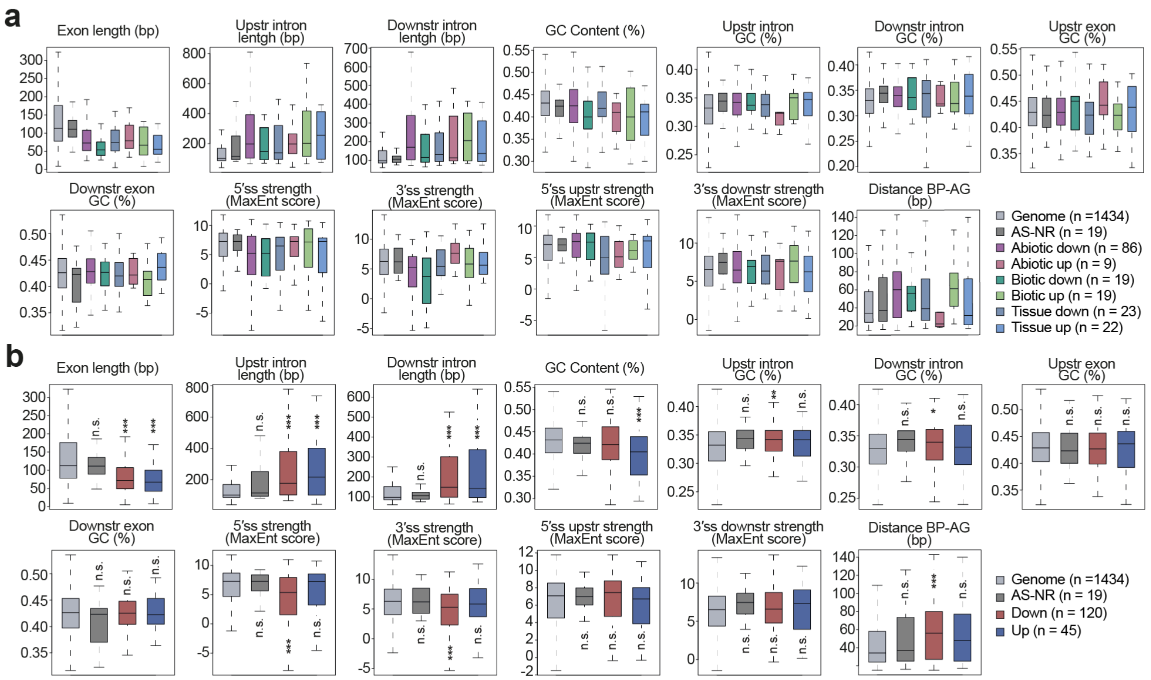


**Figure S18. Genomic regulatory features of differentially spliced exons.** Distributions of the length of alternatively spliced exons together with their downstream and upstream exons, as well as their GC content, maximum entropy (MaxEnt) score for 5′ and 3′ splice sites, and the length between the branch point to the 3′ splice site (BP-AG) for up- and down-regulated exons for each core set (**a**) or for all core sets together (**b**). Genomic features were extracted using Matt [9]. "Genome" contains exons fulfilling the read coverage criteria used for AS analyses. *p*-values indicate statistical significance as assessed by Wilcoxon Rank-Sum tests (P < 0.05 (*); P < 0.01 (**); P < 0.001 (***)). AS-NR, alternative spliced-non regulated; n.s., not significant; Ups: upstream; Downs: downstream.

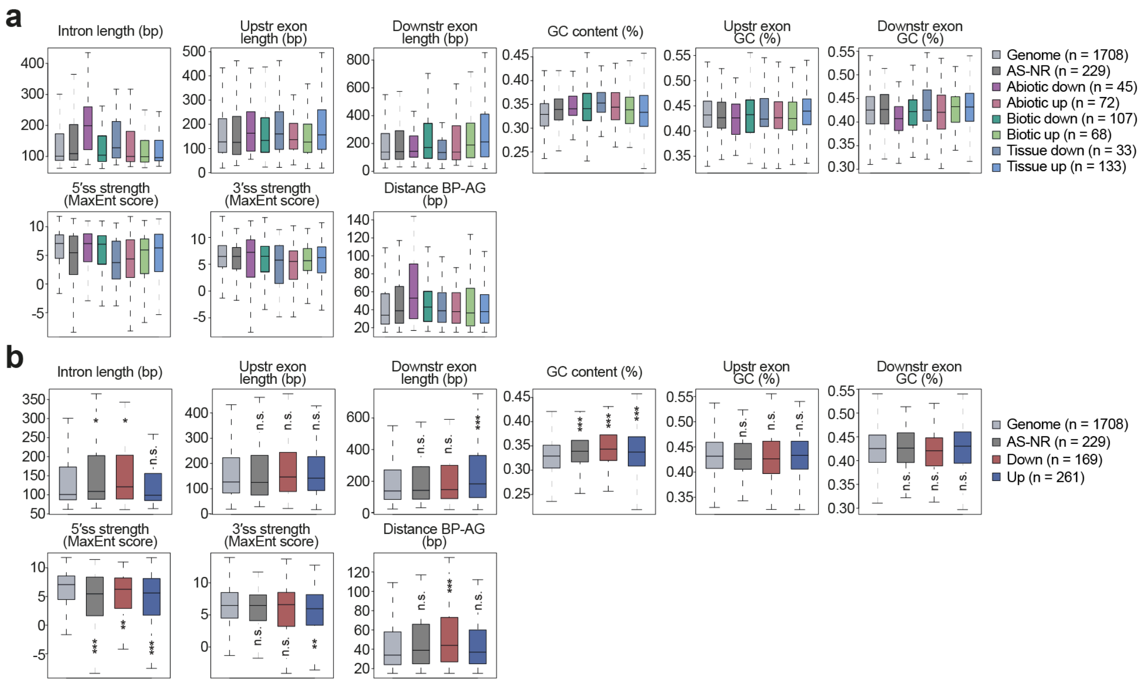


**Figure S19. Genomic regulatory features of differentially spliced introns.** Distributions of the length of alternatively spliced introns together with their downstream and upstream exons, as well as their GC content, maximum entropy (MaxEnt) scores for 5′ and 3′ splice sites, and the length between the branch point to the 3′ splice site (BP) for up- and down-regulated introns from each core set (**a**) or for all core sets together (**b**). Genomic features were extracted using Matt [9]. "Genome" contains introns fulfilling the read coverage criteria used for AS analyses. *p*-values indicate statistical significance as assessed by Wilcoxon Rank-Sum tests (P < 0.05 (*); P < 0.01 (**); P < 0.001 (***)). AS-NR, alternative spliced-non regulated; n.s., not significant.

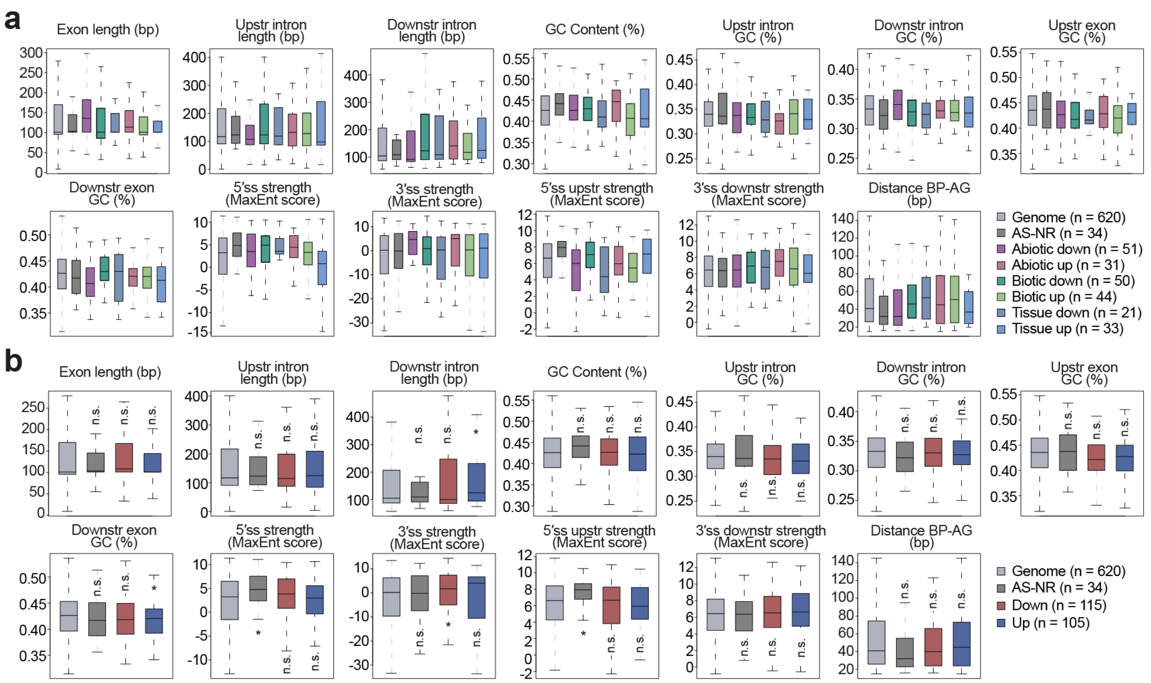


**Figure S20. Genomic regulatory features of differentially spliced alternative splice donors (ALTD).** Distributions of the length of exons containing alternatively spliced donors together with their downstream and upstream exons, as well as their GC content, maximum entropy (MaxEnt) scores for 5′ and 3′ splice sites, and the length between the branch point to the 3′ splice site (BP) for up- and down-regulated splice donors from each core set (**a**) or for all core sets together (**b**). Genomic features were extracted using Matt [9]. "Genome" contains exons associated with ALTD events fulfilling the read coverage criteria used for AS analyses. *p*-values indicate statistical significance as assessed by Wilcoxon Rank-Sum tests (P < 0.05 (*); P < 0.01 (**); P < 0.001 (***)). AS-NR, alternative spliced-non regulated; n.s., not significant; Ups: upstream; Downs: downstream.

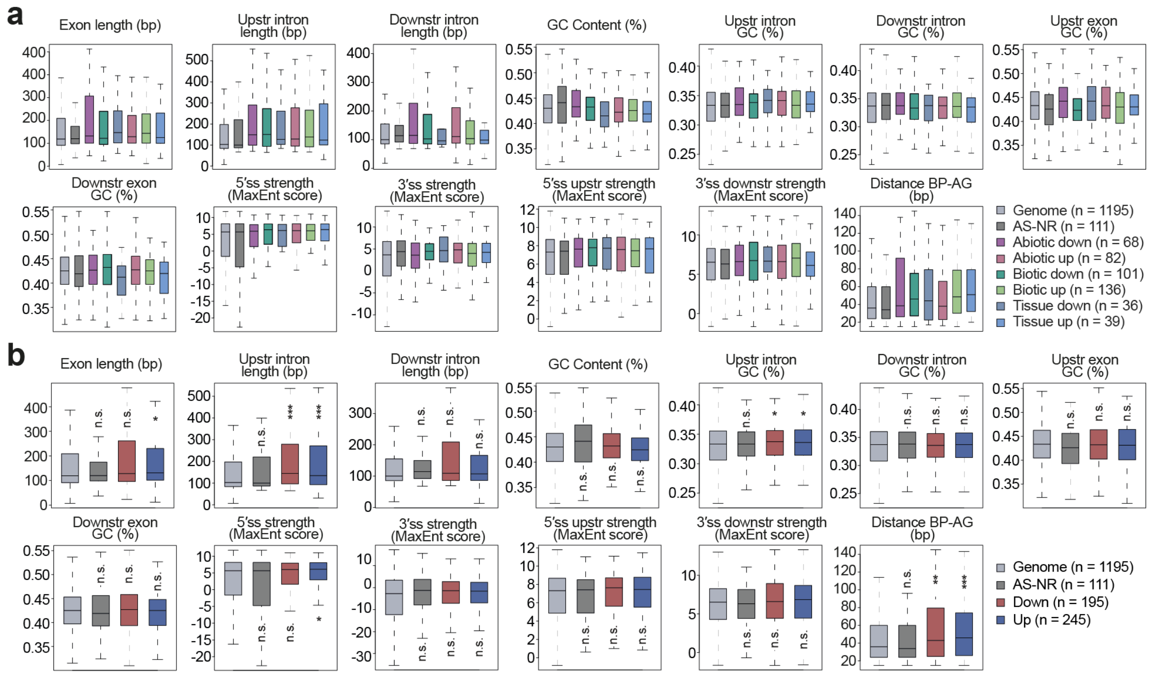


**Figure S21. Genomic regulatory features of differentially spliced alternative splice acceptors (ALTA).** Distributions of the length of exons containing alternatively spliced acceptors together with their downstream and upstream exons, as well as their GC content, maximum entropy (MaxEnt) scores for 5′ and 3′ splice sites, and the length between the branch point to the 3′ splice site (BP) for up- and down-regulated splice acceptors from each core set (**a**) or for all core sets together (**b**). Genomic features were extracted using Matt [9]. "Genome" contains exons associated with ALTA events fulfilling the read coverage criteria used for AS analyses. *p*-values indicate statistical significance as assessed by Wilcoxon Rank-Sum tests ($P < 0.05$ (*); $P < 0.01$ (**); $P < 0.001$ (***)). AS-NR, alternative spliced-non regulated; n.s., not significant; Ups: upstream; Downs: downstream.

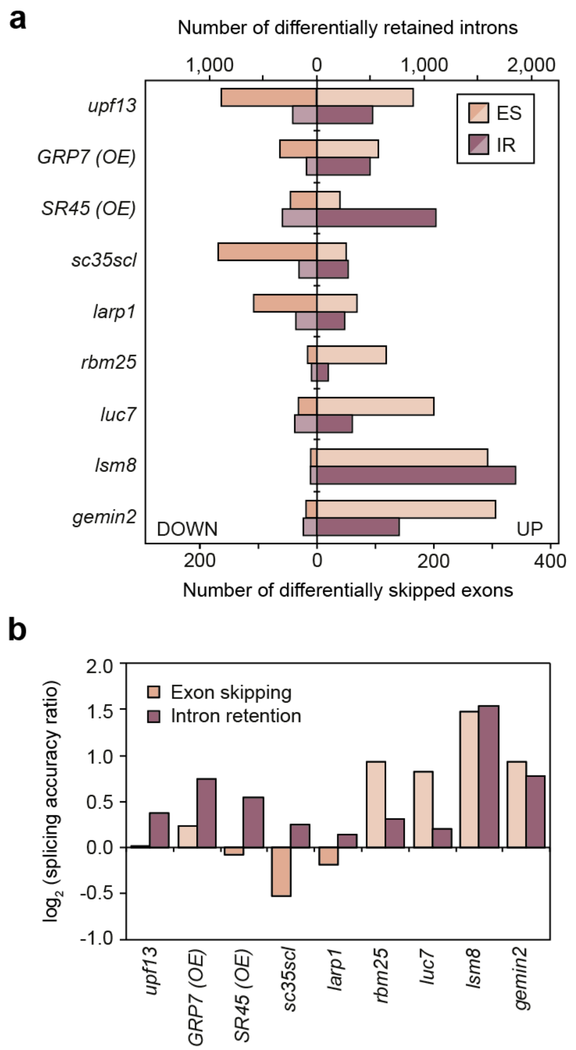


**Figure S22. Global effect of RBP perturbations on ES and IR patterns. a** For each studied RBP experiment, the number of introns with differential retention (top X-axis; right values, higher inclusion in experimental condition) and exons with differential skipping (bottom X-axis; right values, lower inclusion in experimental condition) is shown. **b** Splicing accuracy ratio for each RNA-processing factor mutant. Splicing accuracy ratio is defined, for introns, as the number of introns with increased retention in the experimental condition divided by those with decreased retention, and, for exons, as the number of exons with increased skipping (lower PSI) in the experimental condition divided by those with decreased skipping. The experimental condition corresponds to the respective RBP loss-of-function mutation, with the exception of *gpr7* (overexpression vs. control) and *sr45* (overexpression vs. mutant).

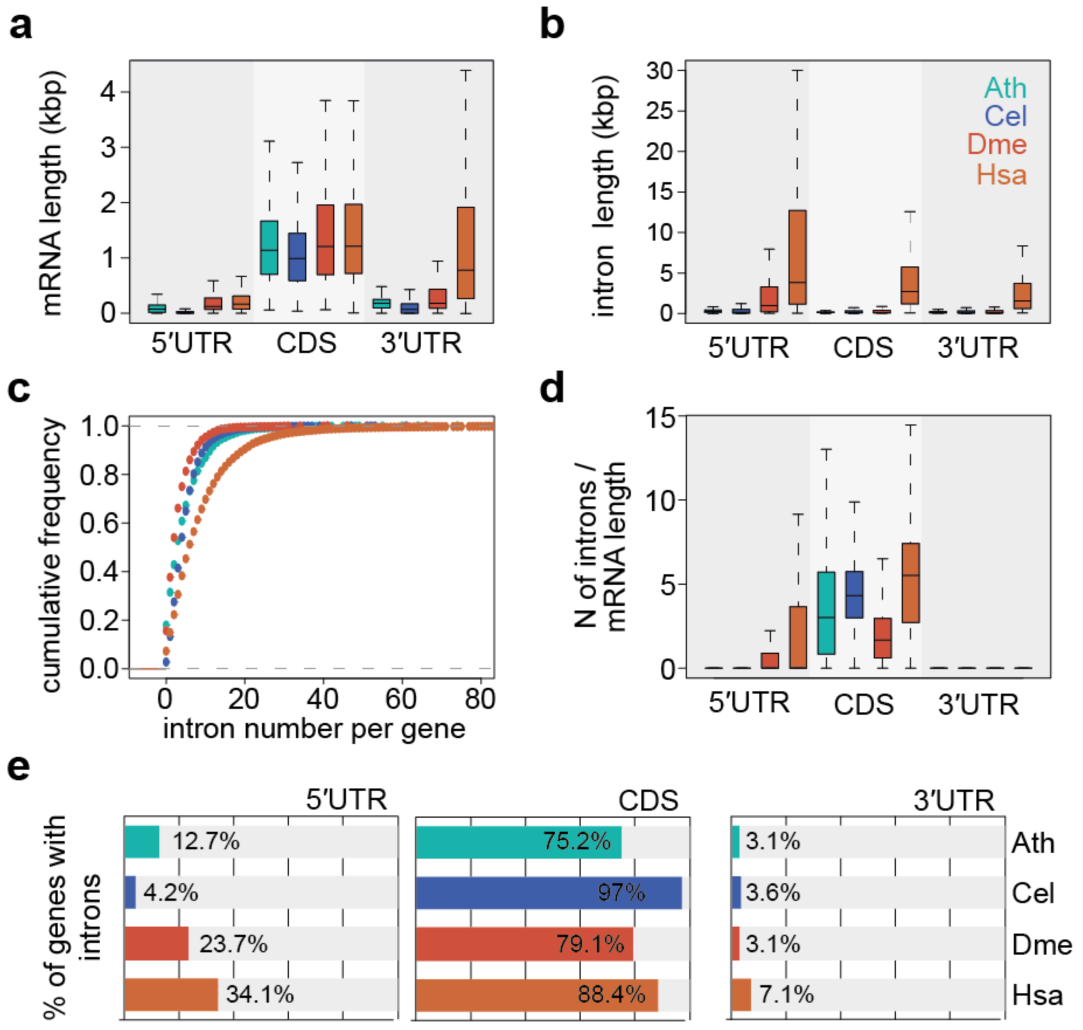


**Figure S23. General genomic features for *A. thaliana*, *C. elegans*, *D. melanogaster* and *H. sapiens*.** Distributions of (**a**) mRNA and (**b**) intron lengths. **c** Cumulative frequency of the number of introns per gene. **d** Intron density. **e** Percentage of genes with introns in the 5′ UTR (left), CDS (middle) and 3′ UTR (right). N, number; CDS, coding sequence; Ath, *A. thaliana*; Cel, *C. elegans*; Dme, *D. melanogaster*; Hsa, *H. sapiens*.

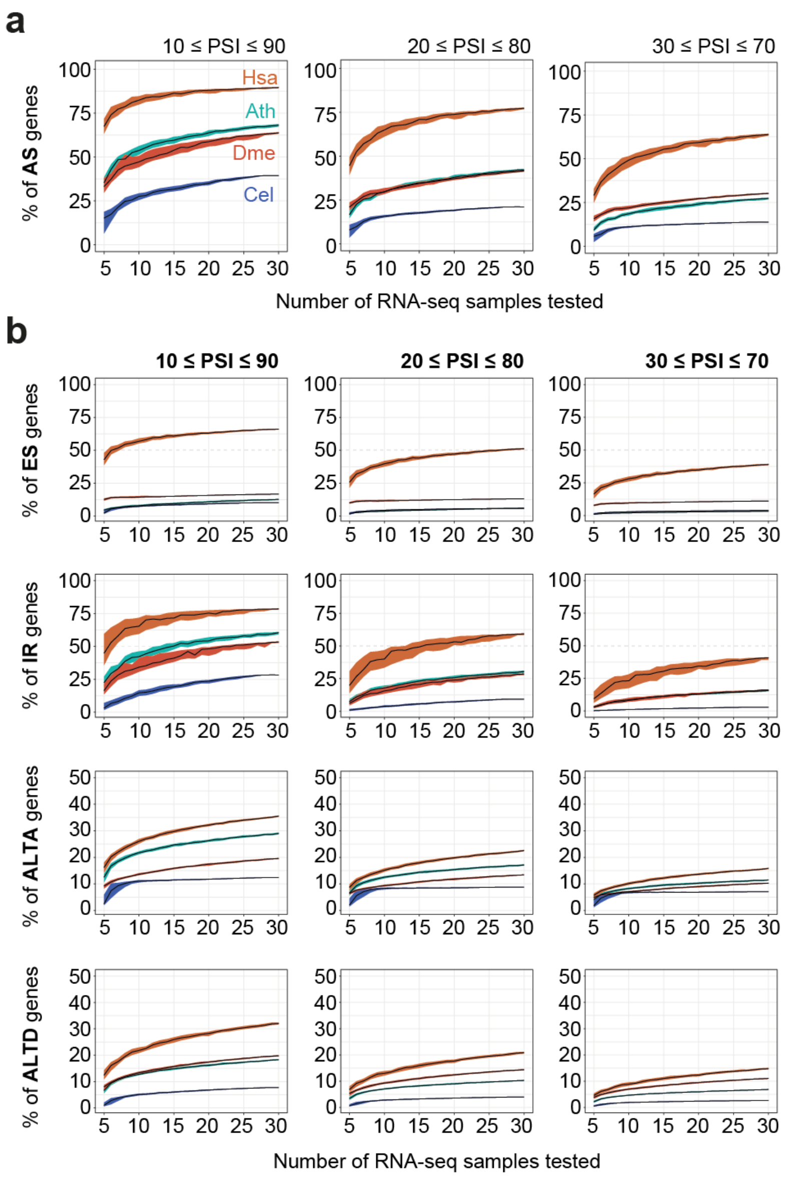


**Figure S24. AS quantifications in *A. thaliana*, *C. elegans*, *D. melanogaster* and *H. sapiens*.** Percent of genes with at least one AS event of all types (**a**) or individual types (**b**) with 10 ≤ PSI ≤ 90 (left), 20 ≤ PSI ≤ 80 (middle) or 30 ≤ PSI ≤ 70 (right) using an increasing number of random samples selected from those used to define the stress and tissue-specific core sets. Only multi-exonic genes with at least one AS event of the tested type(s) with sufficient read coverage in at least 5 samples are considered in each iteration. Dispersion range corresponds to the mean and interquartile range for 100 iterations. ALTD, alternative splice donor; ALTA, alternative splice acceptor; ES, exon skipping; IR, intron retention. Ath, *A. thaliana* (green); Cel, *C. elegans* (blue); Dme, *D. melanogaster* (red); Hsa, *H. sapiens* (orange).

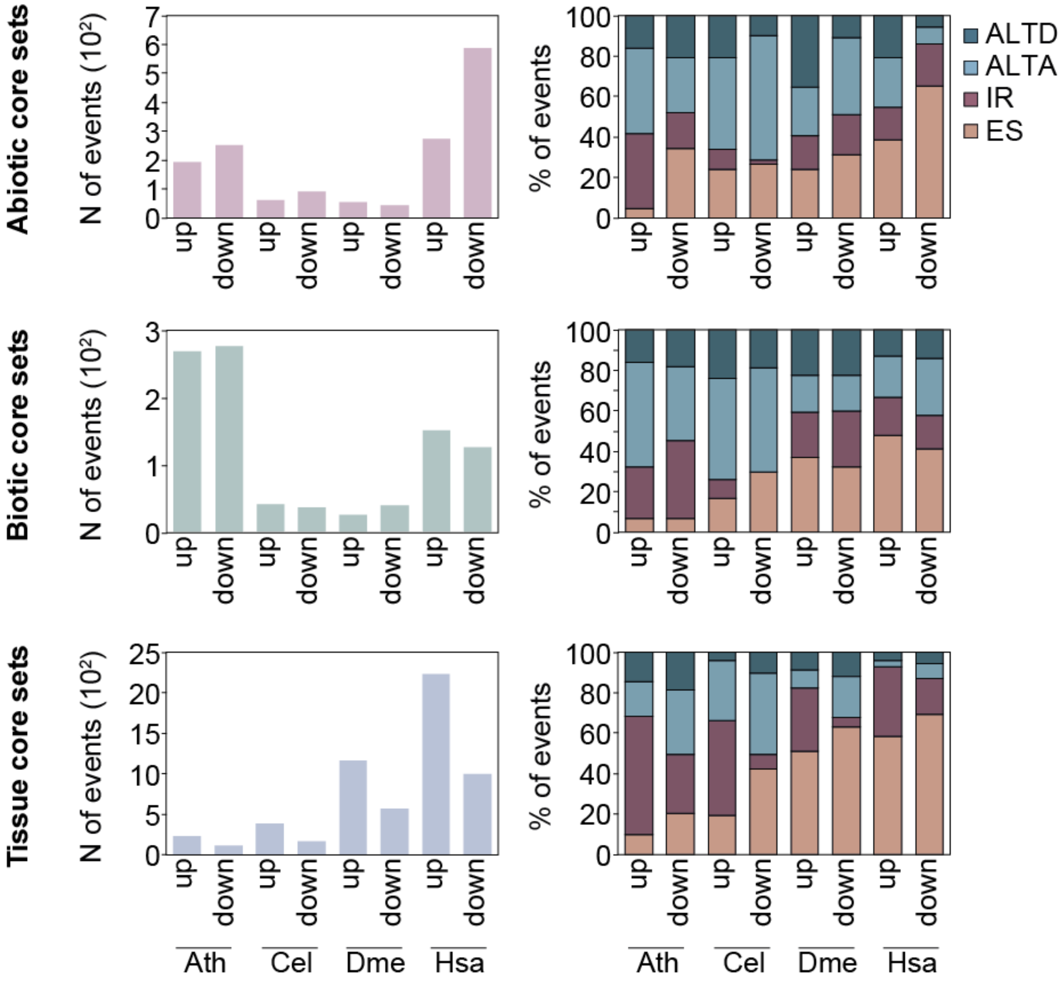


**Figure S25. Number and distribution by type of AS events of each core set in each organism.** Total number of AS events (left) and distribution of AS types (right) for differentially up- or down-regulated AS events belonging to the abiotic (up), biotic (middle) and tissue (bottom) core sets in *A. thaliana* (Ath), *C. elegans* (Cel), *D. melanogaster* (Dme) and *H. sapiens* (Hsa). N, number; ALTD, alternative splice donor; ALTA, alternative splice acceptor; ES, exon skipping; IR, intron retention.

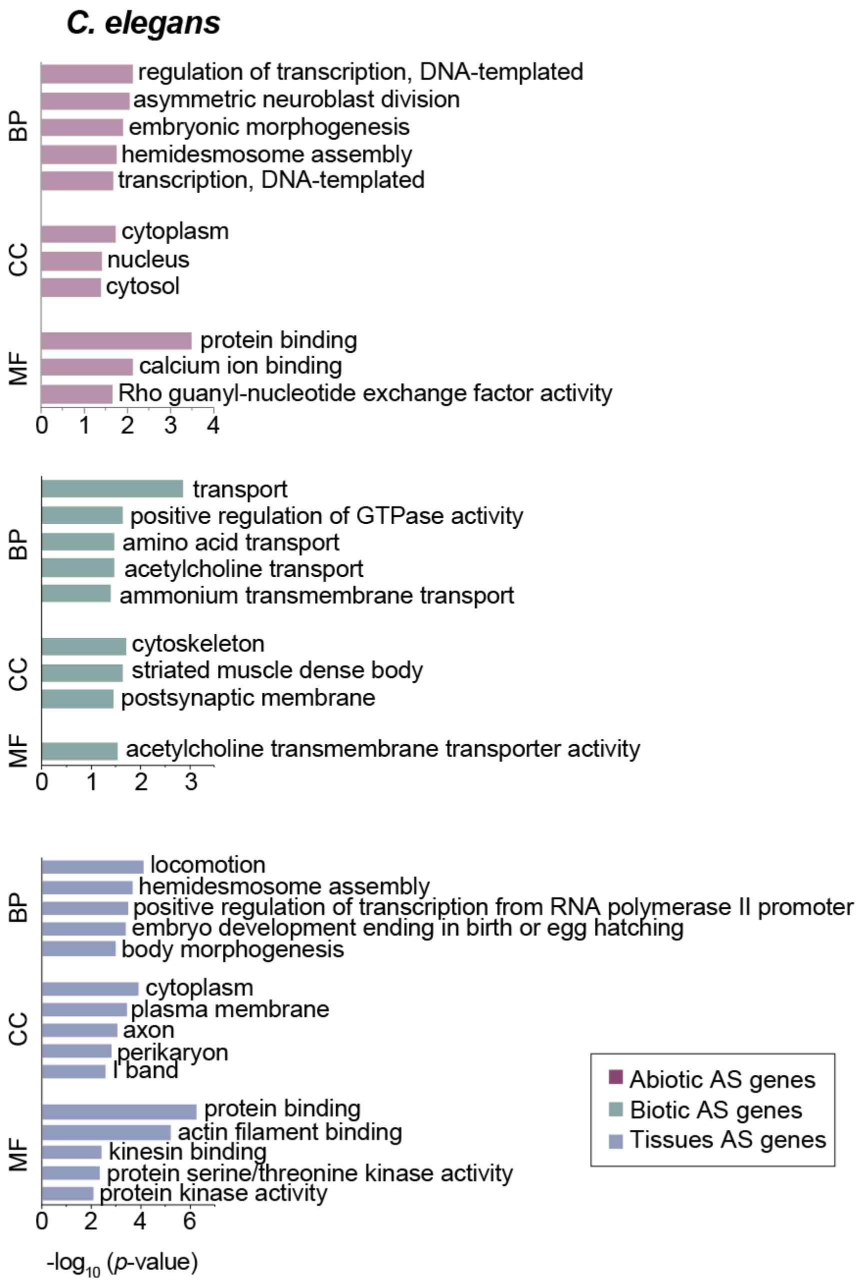


**Figure S26. Functional enrichment analysis of differentially spliced genes in *C. elegans*.** Top five most enriched Gene Ontology (GO) categories for biological process (BP), cellular component (CC) and molecular function (MF) of genes from the abiotic (top), biotic (middle) and tissue (bottom) core sets. $p$-values (Fisher's Exact test) were obtained from DAVID.

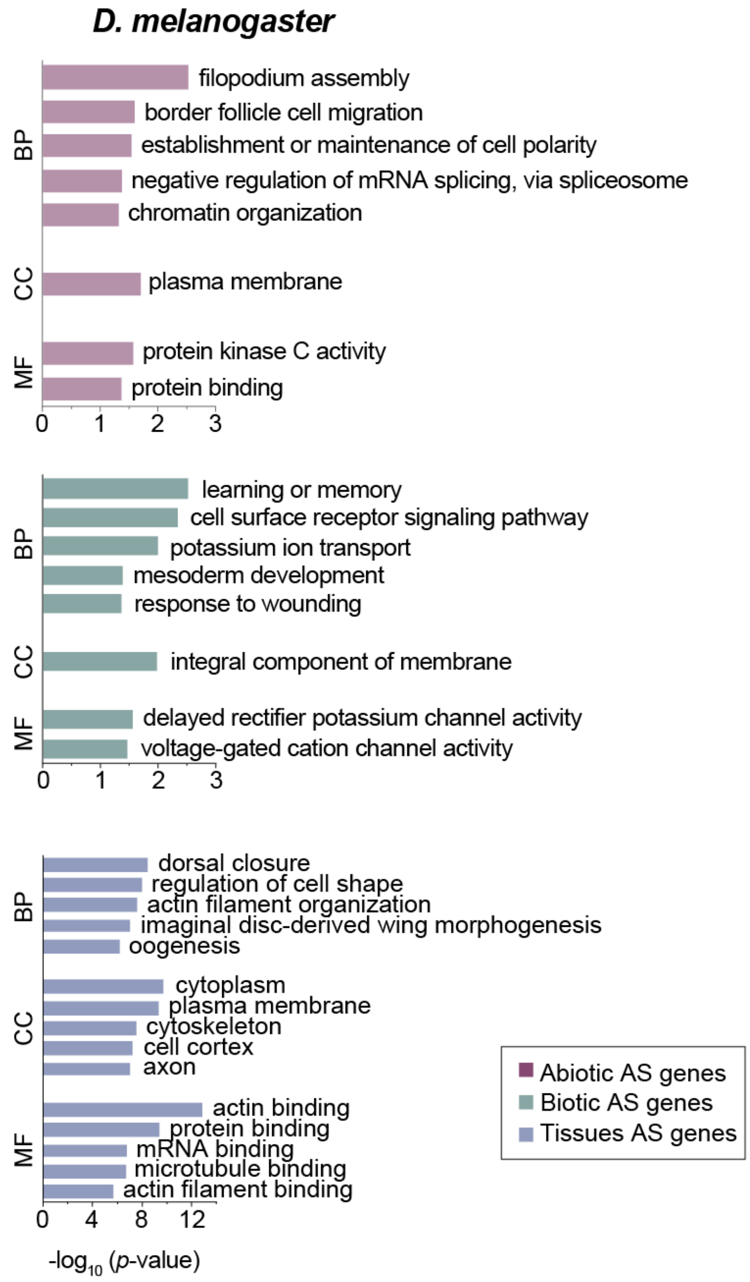


**Figure S27. Functional enrichment analysis of differentially spliced genes in *D. melanogaster*.** Top five most enriched Gene Ontology (GO) categories for biological process (BP), cellular component (CC) and molecular function (MF) of genes from the abiotic (top), biotic (middle) and tissue (bottom) core sets. *p*-values (Fisher's Exact test) were obtained from DAVID.

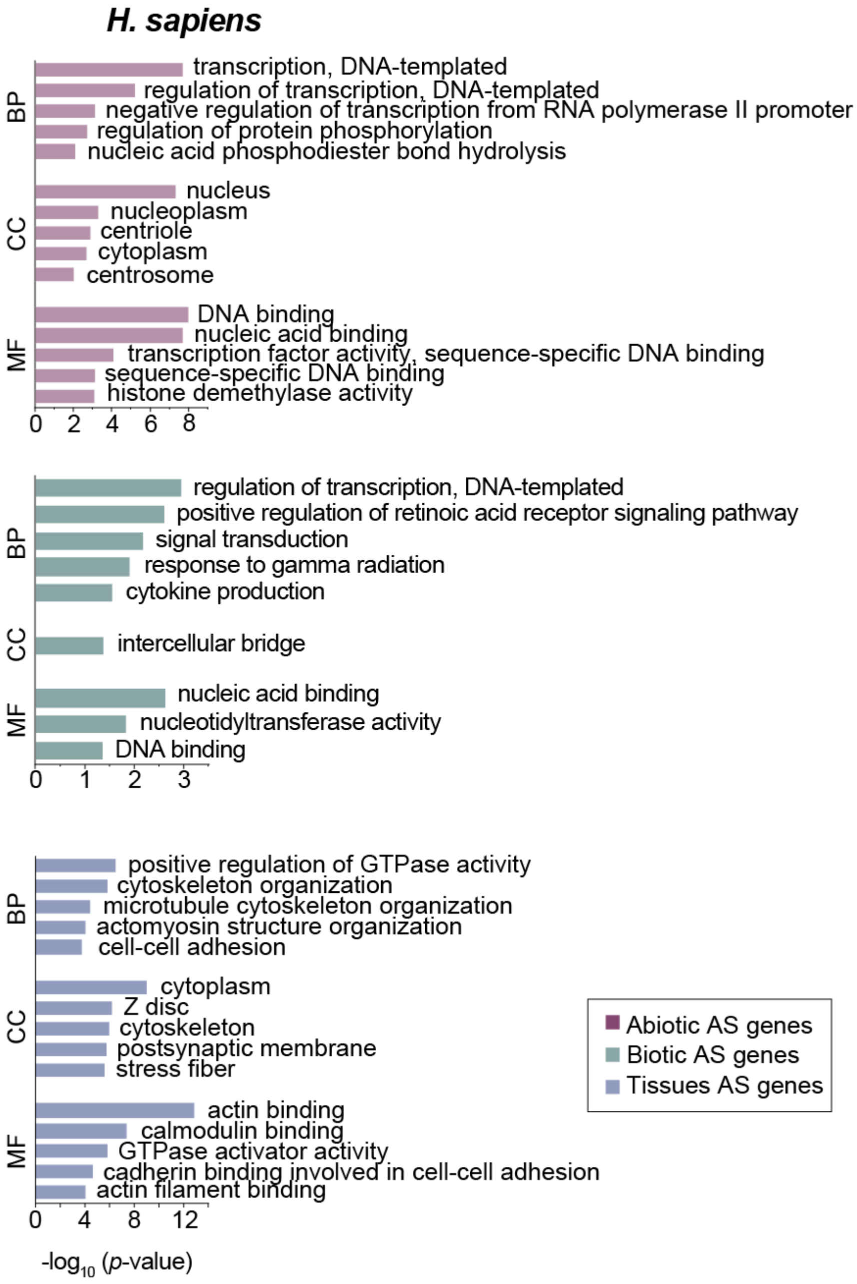


**Figure S28. Functional enrichment analysis of differentially spliced genes in *H. sapiens*.** Top five most enriched Gene Ontology (GO) categories for biological process (BP), cellular component (CC) and molecular function (MF) of genes from the abiotic (top), biotic (middle) and tissue (bottom) core sets. *p*-values (Fisher's Exact test) were obtained from DAVID.

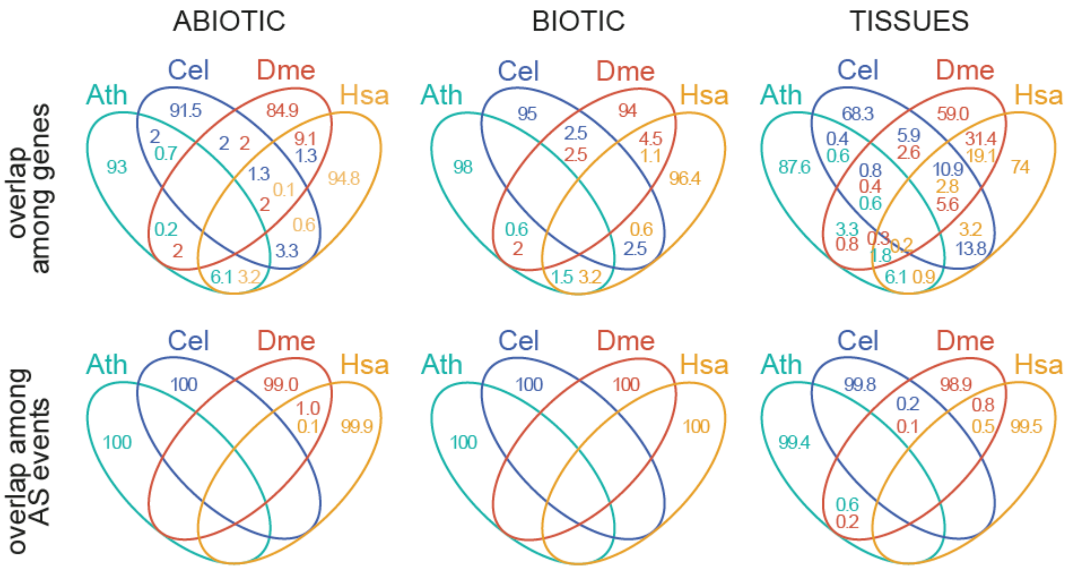


**Figure S29. Comparison of AS core sets across studied species.** Four-way Venn diagrams showing the overlap among AS core sets from *A. thaliana* (Ath), *C. elegans* (Cel), *D. melanogaster* (Dme) and *H. sapiens* (Hsa) at the orthologous gene level (top) and AS event level (bottom). For the AS event level, AS events of any type affecting the orthologous splice site in two different species are considered as overlapping. Note that overlap at either level does not imply evolutionary conservation; at such large evolutionary distances, the overlap is more likely the result of convergent evolution.

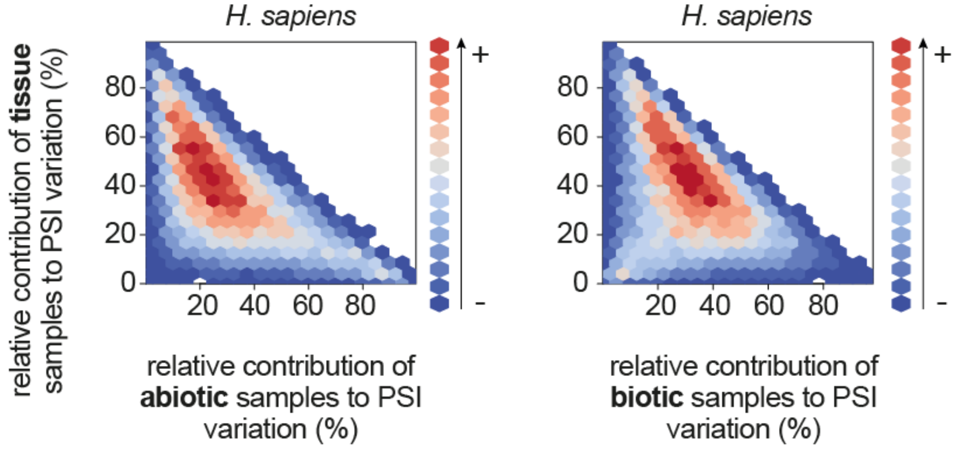


**Figure S30. Relative contribution of tissue and stress samples to PSI variation in *H. sapiens*.** Comparison of the relative contribution to the total PSI variation of tissues used to define the "non-classic" tissue-specific core set vs. abiotic (left) or biotic (right) stress samples in human. The "non-classic" tissue-specific core set was obtained by comparing the following tissues: adipose, colon, lung, ovary and skin. The total PSI variation for each AS event is calculated as the sum of the three relative contributions: (i) the PSI range across tissues, (ii) the maximum ΔPSI among the abiotic experiments, and (iii) the maximum ΔPSI among the biotic experiments (see Methods for details). The color code represents the number of AS events found in each possible intersection between the relative contributions (in percentage) for each set of samples.

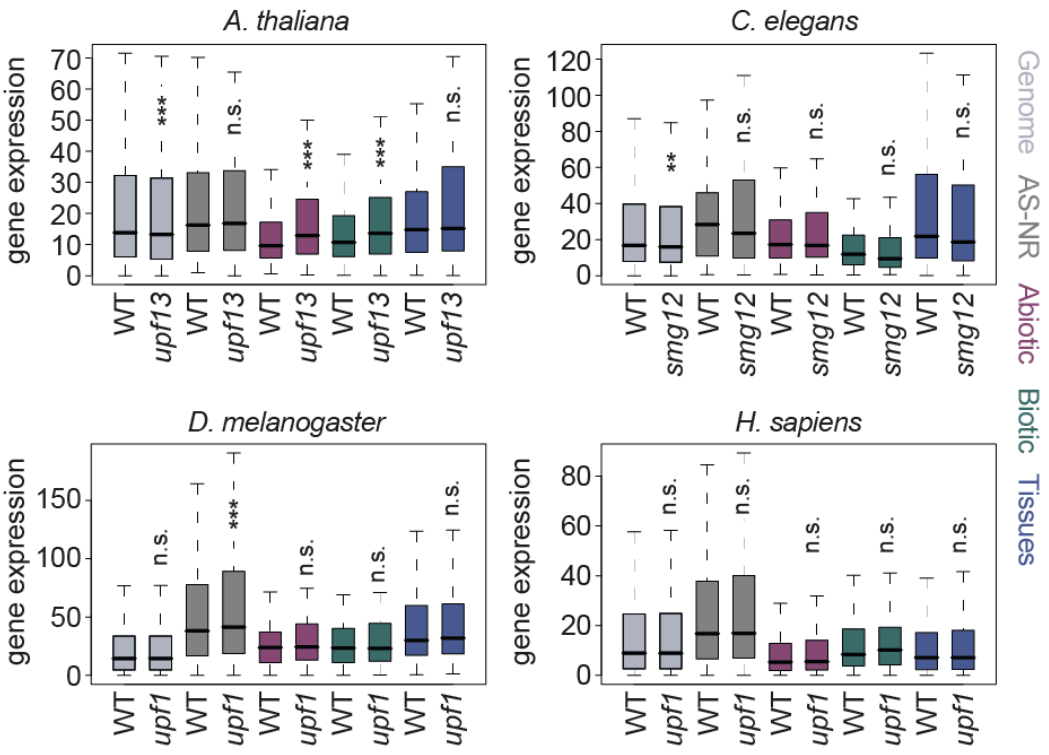


**Figure S31. NMD-regulation of the genes differentially spliced from each core in the different organisms.** Expression values (in cRPKMs) in control and NMD-depleted conditions of genes belonging to the different AS core sets for each species. *p*-values indicate statistical significance as evaluated by Wilcoxon Rank-Sum tests (P < 0.05 (*); P < 0.01 (**); P < 0.001 (***)). "Genome" contains all multi-exonic genes in each species fulfilling the coverage criteria used for the corresponding AS analyses. RNA-seq data was obtained from: *A. thaliana*, [7]; *C. elegans*, [10]; *D. melanogaster*, [11]; *H. sapiens*, [12]. AS-NR, alternative spliced-non regulated; n.s., not significant.

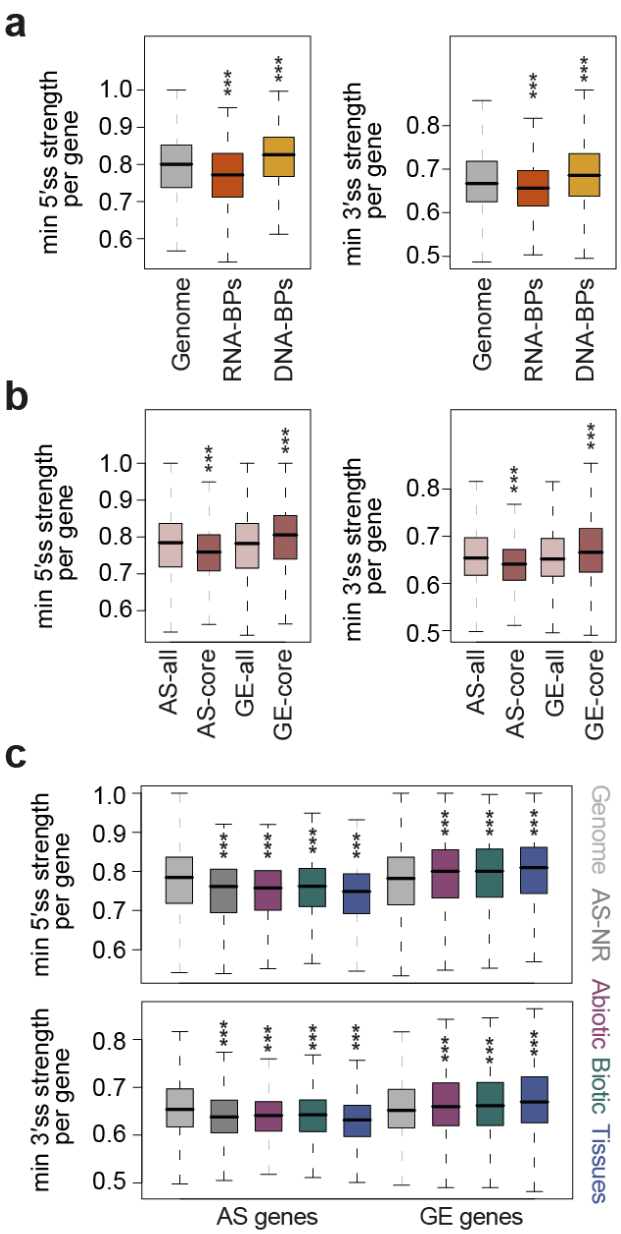


**Figure S32. Comparison of 5' and 3' splice site strengths using an alternative metric.** For all plots, splice site strength was calculated as the similarity of the splice site sequence to the corresponding position-weighted matrix (PWM) built using all annotated 5′ or 3′ splice sites in *A. thaliana*. **a** Distributions of the lowest 5′ splice site (left) or 3′ splice site (right) strength per gene for each category. **b,c** Distributions of the lowest 5′ splice site (**b**, left; **c**, top) or 3′ splice site (**b**, right; **c**, bottom) strength per gene for all AS or GE core set genes together (**b**) or for each AS and GE core set separately (**c**) and matched control sets. Values for all panels correspond to the weakest splice site in each gene.
